# Cellular interactions in the sentinel lymph node predict melanoma recurrence

**DOI:** 10.64898/2025.12.15.694104

**Authors:** Sabrina M. Solis, Yang Yang, Yee Hoon Foong, Anushka Dheer, Kevin L. Ma, Mohammad Saad Farooq, Shira G. Rosenberg, Sophie L. Gray-Gaillard, Giorgos C. Karakousis, Xiaowei Xu, Rami S. Vanguri, Alexander C. Huang, Ramin Sedaghat Herati

## Abstract

Melanoma outcomes have dramatically improved over the past decade, but some patients still experience disease recurrence, particularly those who present at later stage of disease. Prior studies identified an altered immune microenvironment in the sentinel lymph node (SLN) including the presence of dysfunctional CD8 T cells and CD4 T regulatory cells (Tregs), but the spatial organization of the SLN and the interactions of individual cell types have not been extensively studied. To understand how spatial organization of immune cells in the SLN is related to outcomes, we performed spatial proteomic profiling of Stage I and II melanoma SLN to generate an atlas of cell types. Following multiplexed immunofluorescence imaging, deep learning-based segmentation and clustering, we identified 33 subsets of T cells, B cells and Tumor cells. Lymphoid region analyses established a foundational spatial map of the melanoma SLN, revealing higher regional frequencies of activated and memory CD4 T cells in Stage I SLN and consistent positioning of T cells at functional spatial locations. To better understand cellular interactions, we evaluated the immediate neighbors around each index cell using Effect Size Interaction mapping (ESI-map), a novel computational toolkit for understanding spatial interactions. Nearest-neighbor analyses across biological replicates revealed rewiring in cell-cell interacting pairs across disease progression and patient outcomes, particularly with respect to exhausted TOX+ CD8 T cells. Stage II patients who experienced disease recurrence had notable enrichment of cellular interactions between Tregs and TOX+ CD8 T cells that was associated with reduced expression of granzyme B and interferon-γ that was most pronounced in exhausted CD8 T cells. Together, these data demonstrate that the specific spatial interactions between Tregs and exhausted CD8 T cells in the SLN provide a novel immune signature for understanding melanoma outcomes.

**One Line Summary:** Treg-CD8 T cell interactions in the sentinel lymph node predict recurrence outcomes in Stage II SLN+ melanoma patients.

## INTRODUCTION

Advanced-stage melanoma was historically associated with poor clinical outcomes,^1^ prior to the development of immune checkpoint inhibitor (ICI) therapies which dramatically improved median overall survival to 71.9 months.^2^ However, not all patients respond to ICI therapies, and 44-51% of advanced-stage melanoma patients experience disease recurrence despite immunotherapy.^3^ Current recurrence risk stratification methods rely on pathologic features like Breslow’s depth, ulceration of the primary tumor and lymph node tumor burden, but these prediction factors have limited accuracy and exclude potential immunological biomarkers.^4^ The difficulty in predicting melanoma recurrence underscores our incomplete understanding of anti-tumor immunity and the factors that inform heterogeneity in clinical response.

The tumor draining lymph nodes (TDLN) are a critical site of immune cell interaction that shapes subsequent immune responses. The TDLN functions as both a reservoir for T cells and an optimal location for the priming and activation of anti-tumor T cell responses.^5,6^ The sentinel lymph node (SLN) is the most proximal lymph node to the primary tumor and serves as one of the first sites of metastasis prior to more widespread dissemination.^7^ SLN are routinely sampled to inform pathologic disease staging but are primarily evaluated for presence of malignant cells.^8–12^ Although the presence of nodal metastases is a hallmark of clinical Stage II SLN+ melanoma, only some patients experience subsequent recurrence. Prior studies evaluating the SLN have revealed a number of striking features. The SLN undergoes progressive immune remodeling, even prior to tumor seeding.^13^ Architectural alterations to the SLN cause impaired lymphocyte migration into the SLN as well as increased afferent lymphatic flow that can introduce immunomodulatory molecules from the tumor.^13^ Additionally, cellular immune modulation is thought to occur through several mechanisms including an influx of CD4 T Regulatory cells (Tregs), decreased function and number of antigen presenting dendritic cells (DCs), and CD8 T cells with limited functional capacity.^14^ A deeper characterization of the timing and nature of these SLN immune changes is therefore critical to better understand the organization and development of effective anti-tumor immunity.

CD8 T cells are critical elements of the anti-tumor immune response, but their functional capacity may be dampened by a variety of factors. Chronic antigen stimulation leads to CD8 T cell exhaustion characterized by reduced effector function and expression of inhibitory molecules.^15–17^ Exhausted CD8 T cells are found in the primary tumor but also in TDLN.^16^ These cells express PD-1 and CTLA-4, driven by the transcription factor TOX.^18–20^ Recent studies show that the TDLN harbors a pool of stem-like progenitor exhausted CD8 T cells,^5^ characterized by high expression of TCF1, high proliferative capacity, and responsiveness to ICI therapy.^5,15,17,21^ In contrast, exhausted CD8 T cells with low TCF1 expression are terminally differentiated, exhibiting poor functionality and low expression of costimulatory molecules.^15,17^ As anti-tumor T cell responses are primed within the TDLN before trafficking to the tumor,^22^ dissecting the subtypes of exhausted CD8 T cells within this lymphoid compartment is critical to understand their differential impact on tumor control. Beyond these intrinsic factors, extrinsic factors can also impair CD8 T cell effector responses, such as immunomodulatory cytokines from the primary tumor or direct modulation from other cells within the tumor microenvironment. Indeed, Treg mediated modulation of CD8 T cells may serve as one the primary mechanisms for reduced anti-tumor immune responses within the SLN.^23–25^ Tregs can also indirectly affect CD8 T cell responses through regulation of other immune cells including CD4 T cells which are needed to maintain CD8 T cell effector function.^26–31^ Therefore, a deeper mechanistic understanding of CD8 T cell functionality and its modulation by the microenvironment within the SLN is critical for informing differences in clinical outcomes.

Although we have limited understanding of how immune cells organize within the tumor microenvironment to generate anti-tumor immune responses, some recent studies have highlighted the importance of understanding spatial dynamics. For example, CD8 T cells in proximity to macrophage dense areas have been identified as predictors of favorable clinical response in melanoma primary tumors.^32,33^ Additionally, cellular neighborhood analyses found better survival was associated with close proximity of T cells to B cells or cytotoxic lymphocytes in Stage IV melanoma.^34^ Moreover, both proximity of PD-1+ cells to PD-L1+ cells and proximity of cytotoxic CD8 T cells to tumor cells in the melanoma primary tumor are associated with better response to ICI therapy.^35,36^ These studies suggest a more complete understanding of anti-tumor immunity requires further investigation into immune cell-cell communication including in sites where immune responses are organized.

Given the need for novel biomarkers that account for the spatial context of immune cells, we characterized the immune landscape of the melanoma SLN by generating a spatial immune cell atlas. Here, we evaluated the spatial immune landscape for SLN taken at the time of primary resection from patients with Stage I or II melanoma and generated an spatial immune atlas of 33 cell subtypes using multiplex immunofluorescence imaging and RNA sequencing. SLN from patients with clinical Stage II SLN+ melanoma and subsequent recurrence had a gene expression signature of impaired T cell function, though there were only subtle differences with respect to cellular subset frequencies overall or by tissue region. To better understand the influence of cell-cell interactions, we interrogated the immediate neighbors around each cell and evaluated pairwise interactions between subsets. Stage II SLN demonstrated rewiring of pairwise interactions particularly those involving Treg subsets and TOX+ exhausted CD8 T cells. Indeed, interactions between exhausted CD8 T cell subsets and ICOS-expressing Tregs were distinctly enriched in the setting of later recurrence and were associated with reduced expression of granzyme B and interferon-γ in the CD8 T cells involved in the interaction. Together, these data highlight the altered immune interaction landscape of SLN in late-stage melanoma and suggest spatial interactions should be considered as a potential biomarker for predicting outcomes.

## RESULTS

### Multiplex Immunofluorescence Imaging of the Melanoma SLN

To characterize the immune-spatial environment of melanoma SLN, we profiled tissue samples (n = 43) from a cohort of clinical Stage II melanoma that underwent SLN biopsy, including 22 that lacked tumor infiltration (SLN-) and 21 that contained tumor micrometastases (SLN+). At the same time, we also analyzed Stage I SLN (n = 8) as a comparison, which would be the closest to a normal lymph node. Samples were randomly selected from archived samples between 2010-2017 at the University of Pennsylvania. The median age of the cohort was 61 and 76% were male. Approximately 43% of clinical stage II melanoma patients experienced subsequent disease recurrence (**Table S1**). Formalin fixed paraffin embedded (FFPE) SLN from each patient were sectioned at 5 µm thickness. Tumor micrometastases within the SLN were identified by a melanoma pathologist using hematoxylin and eosin staining (H&E) (**Supplemental Figure 1A**). From this cohort we selected 10 Stage II SLN that were SLN+ and 5 Stage I SLN for multiplex immunofluorescence imaging using the <u>CO</u>detection by in<u>DEX</u>ing platform, or CODEX (**Figure 1A, Table S2**). CODEX employs oligo-tagged antibodies and cyclic addition and removal of complementary oligo-tagged fluorescent probes to visualize multiple protein markers on a single tissue sample.^37–39^ We optimized the CODEX experimental workflow for our sample cohort, incorporating a high pH antigen retrieval buffer and photobleaching to reduce autofluorescence.^37,40^ We designed a 43-parameter antibody panel to profile the melanoma SLN (**Figure 1A, Table S3, S4**). Antibodies were selected and screened in human melanoma tissue, resulting in a panel that profiles a range of T and B lymphocytes and their signaling pathways, non-immune cells, and melanoma metastases (**Figure 1B, Supplemental Figure 1B**). Following staining and image acquisition of each lymph node, CODEX images were processed per the manufacturer’s pipeline for deconvolution, tile registration, drift compensation and stitching. Next, using the image analysis platform, HALO, images were first qualitatively assessed for distorted or saturated signals that were manually excluded as artifact (**Supplemental Figure 1C**). The total tissue area varied from sample to sample (range 5.91 – 100.15 mm^2^) (**Table S5**). To identify single cells, we next developed a deep learning-based nuclear segmentation algorithm trained on manual annotations of nuclear boundaries defined by DAPI expression (**Supplemental Figure 1D**). Additionally, we trained a deep learning-based user-defined lymphoid region segmentation algorithm to define several geographic regions within the SLN including T/B Mixed, B Cell Dense, T Cell Dense, Germinal Center (GC) and Tumor (**Supplemental Figure 1E**). Tissue areas defined by minimal cellular density (Background) and areas with distorted tissue signal (Artifact) were excluded from downstream analyses. Lymphoid region segmentation was validated based on visual assessment and protein marker expression in which CD3E had highest expression in the T Cell Dense and T/B Mixed regions, CD20 had highest expression in the B Cell Dense and GC regions, and MART1 had highest expression within the Tumor region (**Supplemental Figure 1F, 1G**).

**Figure 1.**
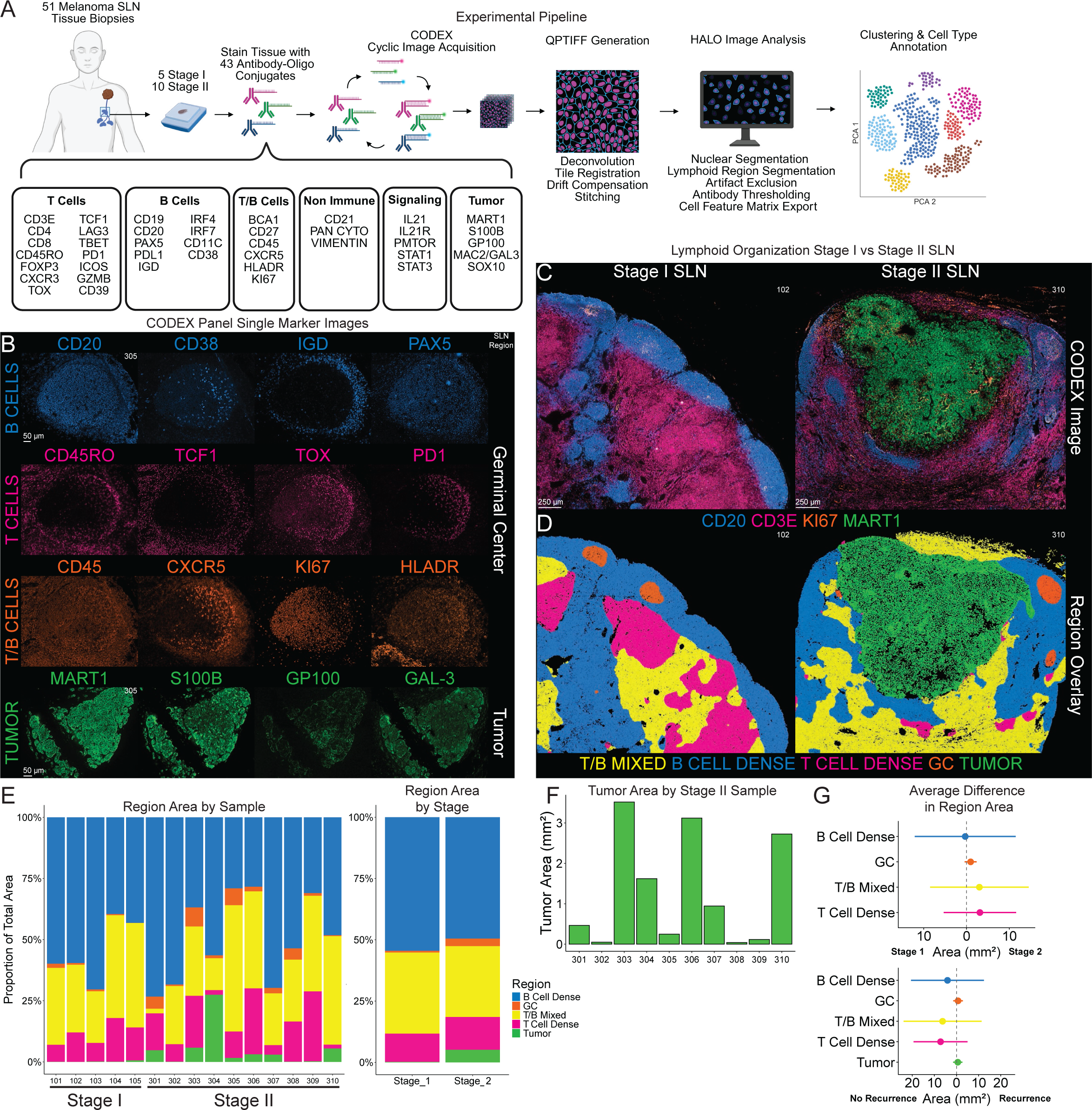
Multiplexed Immunofluorescence Imaging of the Melanoma SLN. **A.** CODEX experimental and analysis workflow. **B.** CODEX antibody panel single marker images from Melanoma SLN tissue with either Germinal Center or Tumor field of view. **C.** CODEX images of Stage I (left) and Stage II (right) Melanoma SLN marking CD20 (blue), CD3E (pink), KI67 (orange), and MART1 (green). **D.** Lymphoid region spatial overlays of corresponding CODEX images shown in 1C, marking T/B Mixed (yellow), B Cell Dense (blue), T Cell Dense (pink), Germinal Center (orange), and Tumor (green) regions. **E.** Area of lymphoid regions as a proportion of tissue area across each SLN sample (left) and averaged by SLN stage (right). **F.** Absolute tumor area across Stage II SLN samples. **G.** Forest plot representing difference in region size by absolute area across Stage (top) and Stage II Recurrence Outcome (bottom).

### Stage II SLN Exhibit Disrupted Lymph Node Architecture

As expected, lymphoid organization was disrupted by the presence of micrometastasis in SLN+ Stage II samples (**Figure 1C**). Whereas Stage I SLN had B Cell Dense regions lining the tissue periphery and T Cell Dense regions were located more centrally, all such structures were disrupted by the infiltration of tumor in Stage II SLN (**Figure 1C, 1D, Supplemental Figure 1H**). Despite this architectural disruption, the average area of lymphoid regions as a proportion of total sample area remained relatively similar across Stage I and Stage II SLN, though individual variation existed (**Figure 1E**). Consistent with clinical observations, micrometastasis size varied across Stage II SLN and were minimally visualized in some samples likely due to the section evaluated (**Figure 1F**).

To determine if these spatial features predicted clinical outcomes, we analyzed absolute lymphoid region area in our cohort, given that six of ten Stage II patients had subsequent recurrence at a median of 10 months after primary resection (whereas the remaining four remained cancer-free after a median of 50.5 months of follow-up). However, we found only subtle differences in area of the T/B Mixed and T Cell Dense regions across stage and outcome (**Figure 1G**) indicating that lymphoid region size alone was not predictive of outcomes in our dataset.

### Immuno-Spatial Characterization Reveals T Cell Alterations in Stage II SLN

To further understand the cellular landscape within the SLN, we quantitatively analyzed our imaging data at the single cell level. The single cell feature matrices (n = 15) were used to generate a Giotto spatial object for each sample (**Figure 2A**).^41^ Following z-score normalization, Giotto objects were integrated using Harmony.^42^ The resulting primary Giotto object contained ∼11.7 million cells across Stage I and II SLN, maintaining information regarding the sample origin and spatial coordinates of the cell (**Figure 2A, Supplemental Figure 2A, 2B**). To annotate cells, we performed high dimensional clustering using the Leiden algorithm.^43^ We identified 33 unique clusters including subsets of CD4 T Cells, CD8 T Cells, B Cells and Tumor Cells (**Figure 2B - 2D, Supplemental Figure 2C**). Cluster classification was validated based on visual assessment of protein marker expression within CODEX images and quantitative evaluation of average marker intensity for each cluster (**Table S6**). We used the x,y coordinates of each cell to computationally generate a spatial projection for a given cluster in each sample. These spatial projections were layered over cells in the CODEX images to validate assigned cluster identities with the cellular phenotypes in the immunofluorescent images (**Figure 2B, 2C**). Clusters that could not be resolved into distinct lineages (i.e. due to expression of CD8 and CD20) were labeled as “Mixed” clusters and retained in the analysis. Some clusters, notably those found within or around the GC, were spatially localized and this was considered when assigning cell type annotations (**Figure 2C**). Overall, this strategy enabled classification of T and B cellular phenotypes within the SLN through integration of high-dimensional clustering with spatial validation using CODEX imaging.

**Figure 2.**
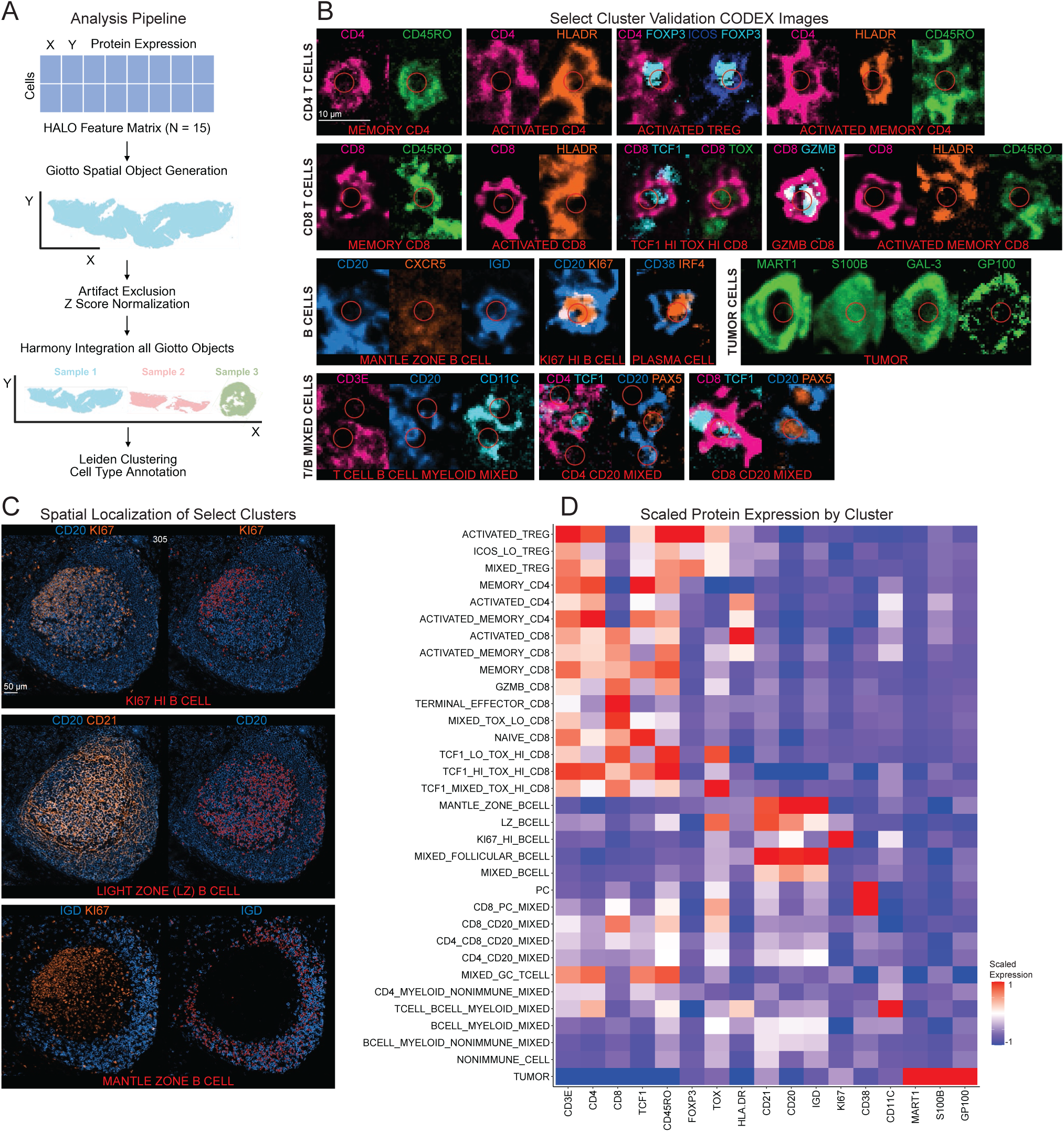
Generating a Cellular Atlas of the Melanoma SLN. **A**. Schematic for Giotto object construction. **B**. Representative CODEX images of cluster phenotypes. Index cell marked by the Red circles. **C**. Representative CODEX images of select cluster phenotypes with spatial localization to the germinal center. **D**. Heatmap for scaled protein expression by cluster.

Most clusters were relatively similar in frequency across Stage I and Stage II SLN and across Stage II recurrence outcomes though individual variation was observed (**Figure 3A, Supplemental Figure 3A**). As expected, however, certain cell types dominated spatial locations within the SLN. For instance, the Germinal Center region was composed primarily of Light Zone B Cells (LZ B CELL), KI67-expressing B Cells (KI67 HI B CELL) and Mixed Follicular B Cells (**Figure 3B**). We next considered whether the subregional composition differed by melanoma stage. Indeed, in the T/B Mixed region, both Activated and Activated Memory CD4 T Cells were more frequent in Stage I SLN compared to Stage II (**Figure 3C**). Similarly, in the T Cell Dense region, Memory CD4 T Cells were more frequent in Stage I compared to Stage II SLN (**Figure 3D**). Conversely, Terminal Effector CD8 T Cells were more frequent in the T Cell Dense region of Stage II SLN compared to Stage I (**Figure 3D**). Although cell subset distribution by region was generally similar, these subtle differences suggested there might be deeper effects on T cell phenotype that warranted transcriptional evaluation.

**Figure 3.**
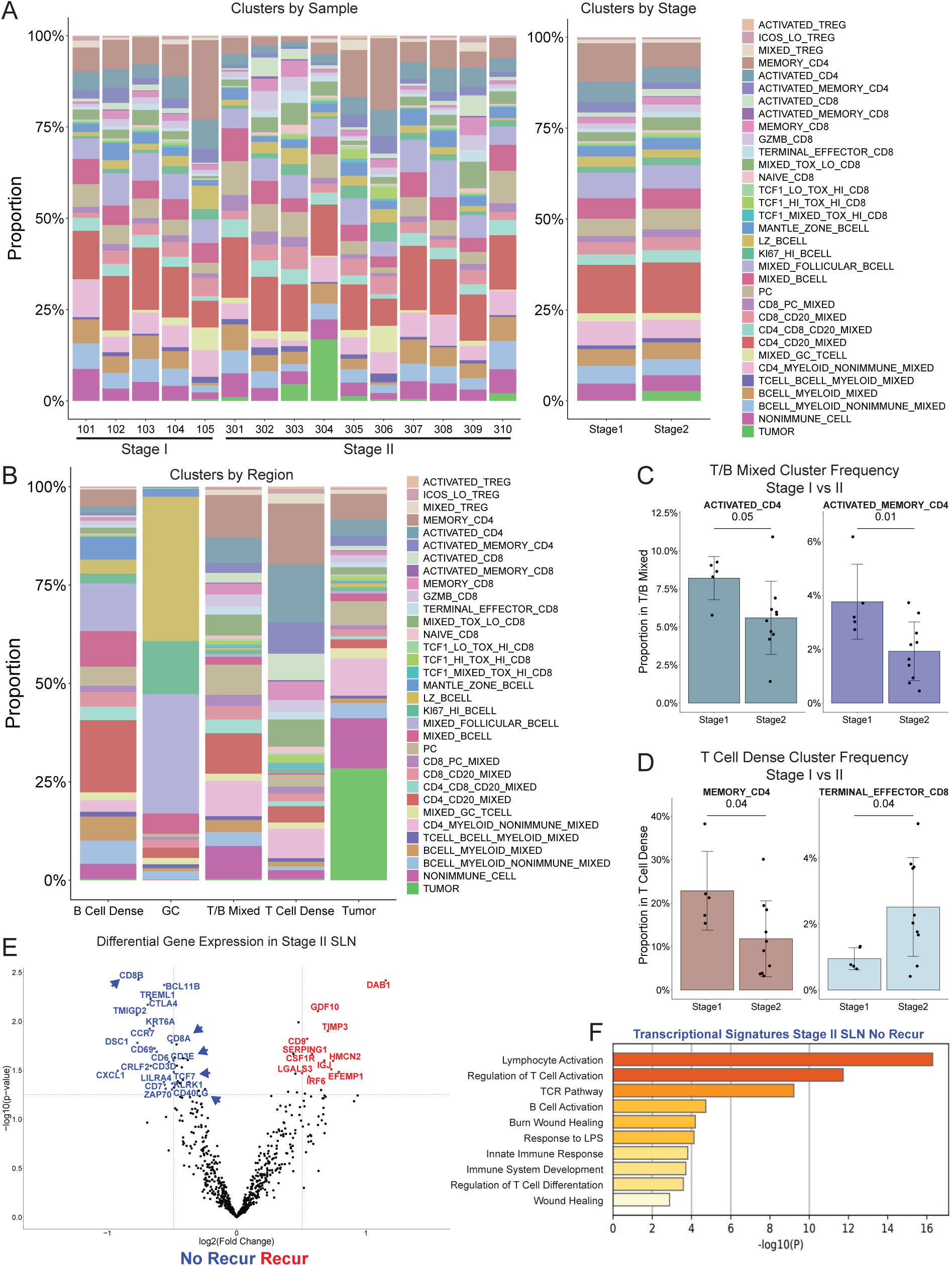
Immuno-Spatial Characterization Reveals T Cell Alterations with Stage II SLN. **A**. Cluster frequencies by sample (left) and averaged across Stage (right). **B**. Cluster frequencies represented as a proportion of cells within each lymphoid region, averaged across all samples. **C-D**. Bar plots displaying region-specific cluster frequency by stage for the frequency of ACTIVATED_CD4 and ACTIVATED_MEMORY_CD4 in the T/B Mixed region (C) and frequency of MEMORY_CD4 and TERMINAL_EFFECTOR_CD8 in the T Cell Dense region (D). P-values determined by unpaired t-test. **E**. Volcano plot from Nanostring analysis representing differential gene expression from Stage II samples excluding micrometastases (blue, No recurrence; red, Recurrence). **F**. Metascape-calculated gene ontology for genes enriched in the No recurrence Stage II SLN signature (Fig. 3E).

Thus, we probed our sample cohort at the transcriptional level with NanoString nCounter gene expression analysis. This technique enables bulk gene expression analysis directly from FFPE tissue samples, using fluorescent probe hybridization and digital counting of mRNA molecules. SLN FFPE tissue samples from Stage I (N = 8), Stage II SLN- (N = 22) and Stage II SLN+ (N = 21) melanoma patients were assessed, including SLN assessed by CODEX. A 3x3 mm margin was defined around any micrometastases for exclusion from Nanostring analysis to ensure profiling of non-involved tissue (**Supplemental Figure 3B**). Analysis of an 800-gene panel revealed that Stage II SLN+ patients who remained recurrence-free had enrichment for *CD3E*, *CD8A*, *CD8B*, *CD40LG* and *TCF7* compared to those that had disease recurrence (**Figure 3E**). The genes enriched in the recurrence-free Stage II SLN+ patients included genes involved in lymphocyte activation and TCR signaling (**Figure 3F**). These data corroborate transcriptional analyses of a study of Stage III melanoma SLN, where recurrence-free patients also had higher expression of genes involved in T cell activation.^44^ Together these findings suggested an association between T cell activation in the SLN and reduced likelihood of disease relapse, thus prompting further focused evaluation of the T cell compartment.

### T Cells Distributed in Immune Active Regions

We next sought to better understand the spatial distribution of CD4 T cells, given the regional enrichment of several activated and memory CD4 T cell populations by frequency within the Stage I SLN and the T cell activation gene signature in Stage II SLN+ patients that remain recurrence-free (**Figure 3**). Using the total number of cells for a given phenotype, we determined cell distribution by calculating the proportion of these cells across our target lymphoid regions. Overall, CD4 T cells were more often present in the T/B Mixed region and less often present in the B Cell Dense region in Stage I SLN compared to Stage II (**Figure 4A**). Next focusing on the activated and memory CD4 phenotypes, these subsets were found in the T/B Mixed and T Cell Dense regions and only rarely were found in Tumor regions (**Supplemental Figure 4A**). Notably, Activated Memory CD4 were more frequently found in the T Cell Dense region relative to Memory CD4 and Activated CD4 (**Figure 4B, 4C**). However, these subsets did not clearly differ in distribution with respect to stage and outcome (**Supplemental Figure 4B, 4C**).

**Figure 4.**
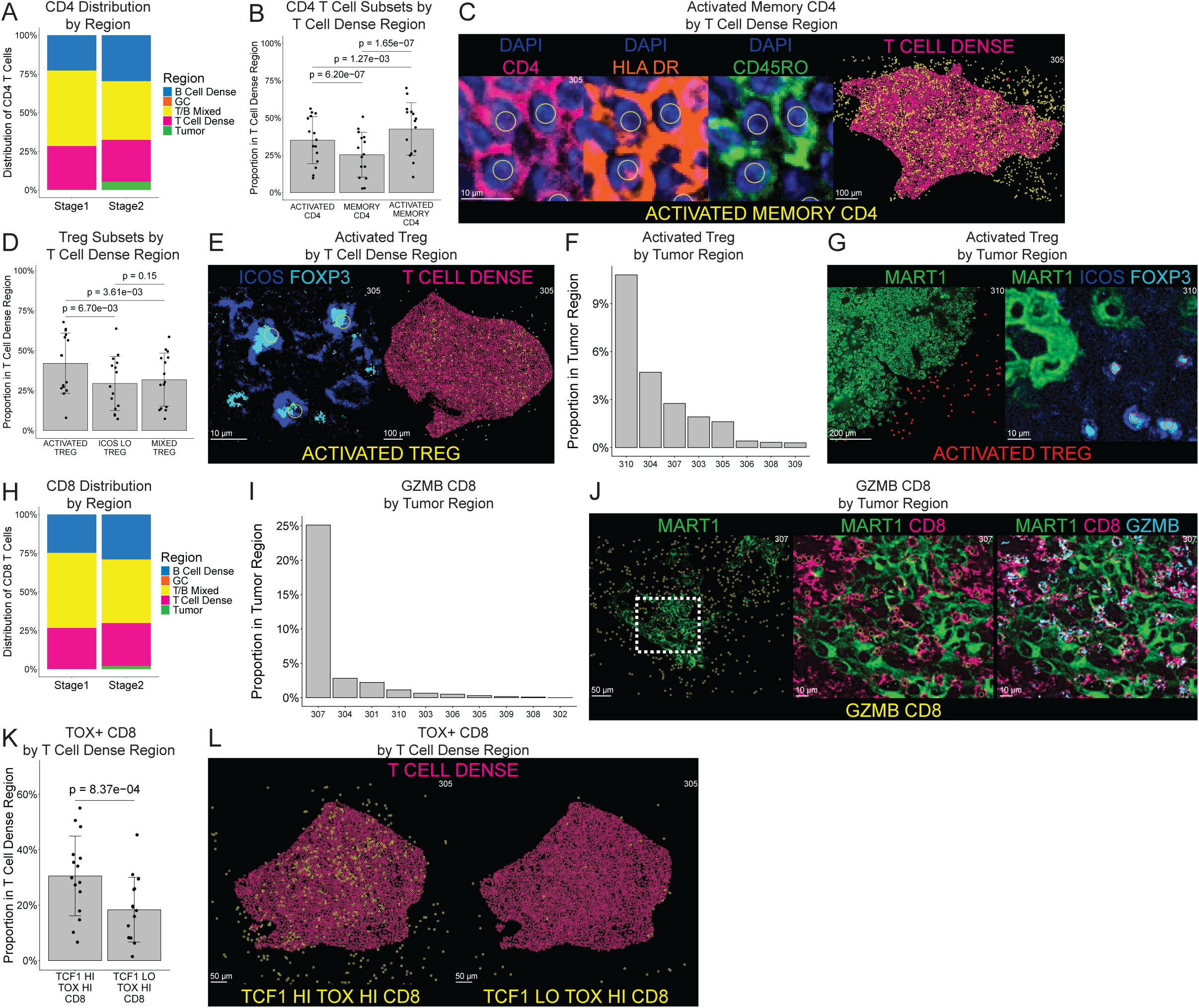
T Cells Distributed in Immune Active Regions in Melanoma SLN. **A**. Stacked bar plot for CD4 T cell distribution by region across SLN stage. **B**. Bar plot comparing proportion of ACTIVATED CD4, MEMORY CD4 and ACTIVATED MEMORY CD4 within T Cell Dense Region. P-values by paired t-test. **C**. CODEX images (left) marking Activated Memory CD4 T cells (yellow circles), DAPI (blue), CD4 (pink), HLADR (orange) and CD45RO (green). Representative image of T Cell Dense region overlay (right, pink) showing Activated Memory CD4 T cell (yellow circles). **D**. Bar plot comparing proportion of ACTIVATED TREG, ICOS LO TREG and MIXED TREG within T Cell Dense Region. P-values by paired t-test. **E**. CODEX image (left) marking Activated Tregs (yellow circles), FOXP3 (cyan) and ICOS (blue). Representative image of T Cell Dense region overlay (right, pink) showing Activated Treg (yellow circles). **F**. Bar plot displaying proportion of ACTIVATED TREG cells found in the Tumor region in the Stage II SLN. **G**. CODEX image marking Activated Tregs (red circles) marked by FOXP3 (cyan) and ICOS (blue) distributed to the tumor micrometastasis defined by MART1 (green). **H**. Stacked bar plot displaying CD8 T cell distribution by region across SLN stage. **I**. Bar plot displaying proportion of GZMB CD8 T cells found in the Tumor region in the Stage II SLN. **J**. CODEX image marking GZMB CD8 T cells (yellow circles) marked by CD8 (pink) and GZMB (cyan) found in micrometastases as defined by MART1 (green). **K**. Bar plot comparing proportion of TCF1 HI TOX HI CD8 and TCF1 LO TOX HI CD8 within the T Cell Dense Region. P-value by paired t-test. **L**. Representative image of T Cell Dense region overlay (pink) displaying TCF1 HI TOX HI CD8 (left, yellow circles) and TCF1 LO TOX HI CD8 (right, yellow circles).

Given prior reports of increased Treg frequency in advanced stage melanoma TDLN,^14,45^ we next considered the spatial positioning of Treg subsets. Treg subsets were identified based on high expression of FOXP3 and stratified by ICOS expression, with high ICOS expression indicating T cell activation and increased suppressive activity (**Figure 2D, Supplemental Figure 4D**).^46^ We identified 3 subtypes of CD4 Tregs: Activated Tregs, ICOS Low Tregs and Mixed Tregs. The Treg subsets in the SLN were primarily found in the T/B Mixed and T Cell Dense regions (**Supplemental Figure 4E**). Activated Tregs had higher cellular proportions in the T Cell Dense region compared to both the ICOS Low and Mixed Tregs (**Figure 4D, 4E**), but their distribution did not differ with respect to stage or recurrence (**Supplemental Figure 4F, 4G**). Varying proportions of Activated Tregs were found near Tumor micrometastases across Stage II SLN (**Figure 4F, 4G**). Overall, these results provide insight to the heterogeneity in location in Treg subsets but did not clearly explain patient outcomes.

As CD8 T cells can help mediate anti-tumor immunity, we next characterized the spatial organization of the CD8 T cell compartment within the SLN. Like CD4 T cells Stage I SLN had more CD8 T cells in the T/B Mixed region compared to Stage II, and Stage II SLN had slightly more CD8 T cells in the B Cell Dense region as well as a fraction of cells within the Tumor region (**Figure 4H**). We first evaluated the 7 unique non-exhausted (TOX Low) CD8 T cell populations: Activated CD8, Activated Memory CD8, Memory CD8, Mixed TOX Low CD8, Naïve CD8, Terminal Effector CD8 and Granzyme B-expressing (GZMB) CD8 T cells. The activated, memory and naïve subsets were primarily found in the T/B Mixed and T Cell Dense regions, and few CD8 T cells were identified within Tumor region, reflecting the trends we observed in the CD4 compartment (**Supplemental Figure 4H, 4I, 4J**). Of note, varying proportions of GZMB CD8 were found within the Tumor region across Stage II SLN but this did not differ by recurrence status (**Figure 4I, 4J, Supplemental Figure 4J**). These data demonstrate that while functional CD8 T cells were observed within immune active regions of the SLN, these patterns were not associated with stage or prevention of recurrence.

We next characterized the exhausted CD8 T cell landscape in the melanoma SLN. We identified 3 exhausted CD8 T cell subsets based on high expression of TOX that were then stratified by TCF1 expression: TCF1+ TOX+ CD8 (Progenitor-Like), TCF1 Mixed TOX+ CD8, and TCF1- TOX+ CD8 (Terminal-Like) (**Figure 2D**). Like other CD8 and CD4 populations, exhausted CD8 T cells were primarily distributed among the T/B Mixed and T Cell Dense regions (**Supplemental Figure 4K**). Notably, TCF1+ exhausted CD8 T cells were more often found in the T Cell Dense region compared to TCF1- exhausted CD8 T cells (**Figure 4K, 4L**). This is consistent with previous reports that Progenitor Exhausted CD8 T Cells are found in the T Cell Zone of uninvolved LNs of head and neck squamous cell carcinoma patients.^47^ Overall, however, there were only subtle differences in spatial distribution of exhausted CD8 T cell subsets based on stage or recurrence (**Supplemental Figure 4L, 4M**).

### Activated Tregs Interact with Exhausted CD8 T Cells in Stage II Recurrence SLN

Spatial profiling of lymphocytes subsets revealed subtle differences in regional distribution, but did not reliably distinguish patients by stage or by recurrence status (**Figure 4**). We reasoned that the direct evaluation of cellular neighborhoods may help clarify mechanisms of T cell impairment, given reduced expression of genes associated with T cell activation in SLN of Stage II SLN+ individuals who did not have recurrence (**Figure 3E, 3F**). To test this idea, we developed Effect Size Interaction mapping (ESI-map), a new toolkit for study of spatial interactions of cellular neighbors across biological replicates.

We first generated a spatial k-Nearest Neighbors (KNN) network to profile the cell-cell interactions in the SLN. For each sample, the most-proximal neighbors (k=9) for each cell were identified (**Supplementary Figure 5A**). For any given cell type, the most frequent neighbors were first calculated as a proportion of the total neighbors for that cell type across all samples. We first validated our approach through evaluation of Light Zone B Cells (LZ B Cells) (**Figure 5A**). As expected, LZ B Cells were most frequently neighbored by other LZ B Cells, as well as other GC cell subtypes including Mixed Follicular B Cells, Mixed GC T Cells, and KI67 HI B Cells (**Figure 5B, Supplementary Figure 5B**).

**Figure 5.**
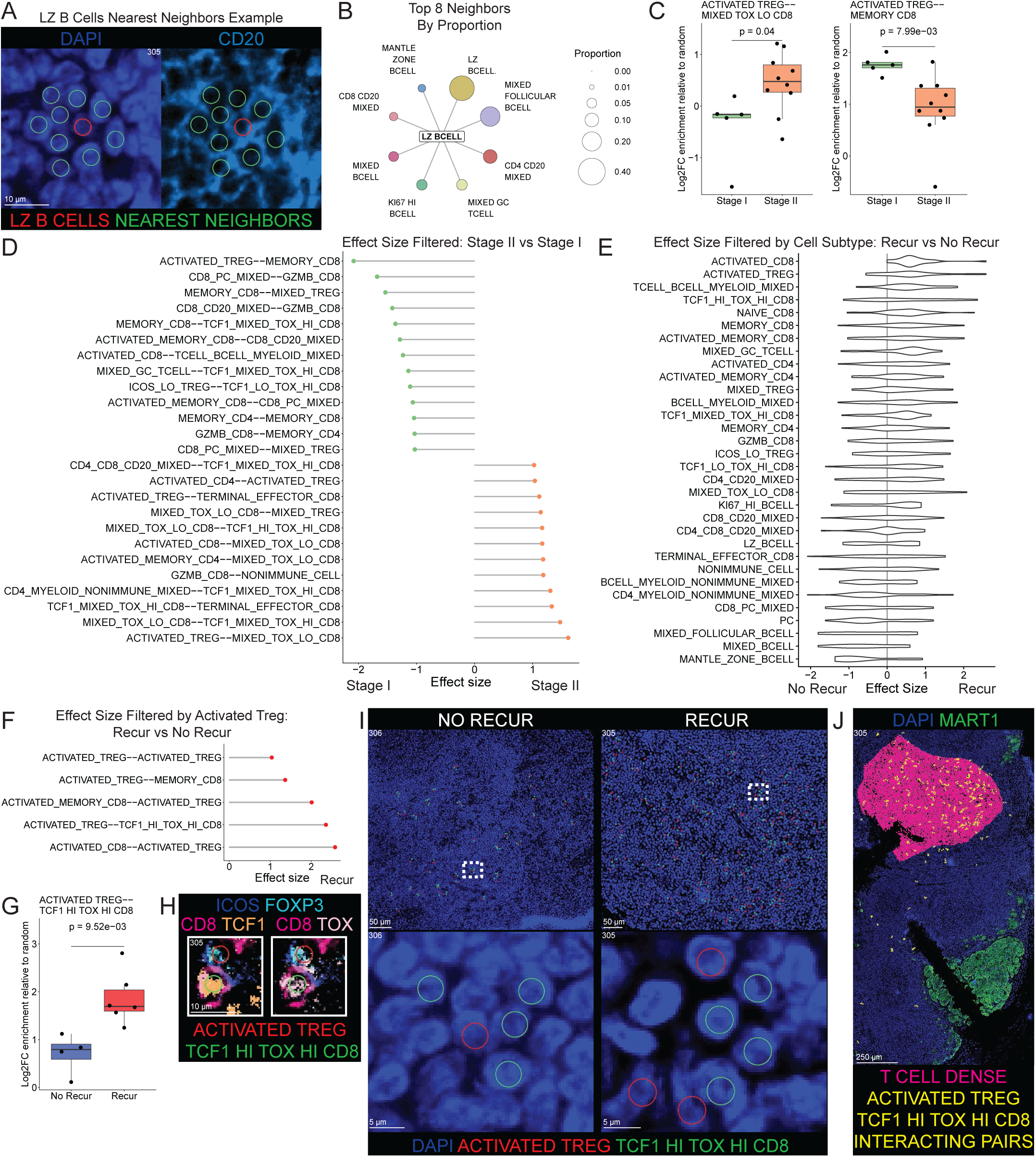
Activated Tregs Interact with Exhausted CD8 T Cells in Stage II Recurrence SLN. **A**. Representative CODEX image of an example LZ BCELL (Index Cell, red) and its nine nearest neighbors (green). Marking CD20 (light blue) for LZ B CELLS and DAPI (royal blue) for all other cells. **B**. Spoke plot displaying top 8 most frequent neighbors of LZ BCELLS by proportion of all LZ BCELL neighbors; size of circle represents proportion amongst neighbors. **C**. Box plot displaying the Log2FC enrichment of interaction pairs relative to random across Stage for ACTIVATED TREG -- MIXED TOX LO CD8 (left) and ACTIVATED TREG -- MEMORY CD8 (right). **D**. Effect size of interaction pairs enriched in Stage II (orange) vs Stage I (green) SLN. Effect size was calculated as the difference in the mean Log2FC enrichment of interaction pairs relative to random, divided by the pooled standard deviation. **E**. Effect size summary plot of cell subtypes involved in interaction pairs enriched in Stage II Recur vs No Recur SLN. **F**. Effect size of interaction pairs involving ACTIVATED TREGS enriched in Stage II Recur (red) vs No Recur SLN. **G**. Box plot displaying the Log2FC enrichment of interaction pairs relative to random for ACTIVATED TREG -- TCF1 HI TOX HI CD8. **H**. Example CODEX image of ACTIVATED TREG -- TCF1 HI TOX HI CD8 interacting pairs; red circle indicates ACTIVATED TREG defined by FOXP3 (cyan) and ICOS (blue) expression, green circle indicates TCF1 HI TOX HI CD8 defined by CD8 (dark pink), TCF1 (orange) and TOX (light pink) expression. **I**. Example CODEX image of ACTIVATED TREG -- TCF1 HI TOX HI CD8 interacting pairs across no recur (left) and recur (right) Stage II SLN tissue. Red circles indicating ACTIVATED TREG, green circles indicating TCF1 HI TOX HI CD8, and DAPI (royal blue) shown. **J**. Representative CODEX image of ACTIVATED TREG -- TCF1 HI TOX HI CD8 interacting pairs (yellow circles), with the T Cell Dense region (pink), MART1 (green) and DAPI (royal blue) shown.

We next compared cell-cell interactions between Stage I and Stage II SLN. We first determined an enrichment score for each possible pairwise interaction as the log_2_ (fold-change) of observed interactions relative to random pairings within the tissue from each individual SLN sample, thereby also accounting for varying cellular frequencies per subset and across patient samples. Next, an effect size was calculated as the difference in the mean enrichment scores across biological replicates for each cohort of interest, divided by the pooled standard deviation. Interrogating the 1089 possible pairwise interactions with this effect size metric revealed a significant rewiring of cellular interactions by stage. Indeed, many cell subtypes involved in pairwise interactions demonstrated large effect sizes (> 1 or < −1) (**Supplementary Figure 5C**), despite the lack of overt differences in subset frequency (**Figure 3, 4**). We then considered specific pairwise interactions identified by the effect size metric. Notably, Activated Tregs had more interactions with Mixed TOX LO CD8 T cells and fewer interactions with Memory CD8 T cells in Stage II compared to Stage I SLN (**Figure 5C, 5D**). Although the top 8 most common neighbors of Activated Tregs were generally similar by frequency in Stage II relative to Stage I SLN, Memory CD8 T cells were notably less frequent neighbors with advanced stage disease (**Supplementary Figure 5D, 5E**). Thus, the spatial interaction landscape of SLN shifted with advanced disease, with particular enrichment of interactions involving Activated Tregs.

To next determine if the changes in cell-cell interactions were associated with outcomes, we focused on recurrence in Stage II SLN+ patients. Effect size was again calculated, now representing the differences in pairwise interactions between Stage II SLN+ patients who remained recurrence free compared to Stage II SLN+ patients who experienced disease recurrence. Compared to the earlier comparison by stage (**Figure 5D**), analysis of recurrence revealed many more pairwise interactions were notable for large effect size (**Supplementary Figure 5F**). Once again, pairwise interactions involving Activated Tregs had large effect sizes associated with recurrence (**Figure 5E**). Indeed, Activated Treg interactions with CD8 T cell subsets were particularly enriched in SLN from patients who later had recurrence (**Figure 5F, Supplemental Figure 5F**). Notably, interactions between TCF1+ TOX+ CD8 T cells and Activated Tregs were enriched in patients who had recurrence (**Figure 5G - 5I**). These interacting pairs were present predominantly in the T Cell Dense region (**Figure 5J, Supplementary Figure 5G**), underscoring previous reports that immunomodulatory phenotypes localize at SLN regions distant from tumor micrometastases.^44^ Furthermore, pairwise interaction enrichment was largely similar across recurrence outcomes whether evaluating via nearest-neighbor approach or by a simple radius (r = 6 µm) approach (**Supplementary Figure 5H**), thereby demonstrating the observation was independent of the method used to identify immediate spatial neighbors. Taken together, these data demonstrate rewiring of cellular interactions with advanced stage disease and highlight the relationship of specific spatial interactions with subsequent recurrence.

### Treg Proximity is associated with reduced effector expression in Exhausted CD8 T Cells

Given the enrichment of Activated Treg interactions with exhausted CD8 T cells in the setting of recurrence, and since ICOS expression on Activated Tregs is associated with increased suppressive activity,^46^ we next considered if the proximity of Treg could functionally influence the protein expression of cellular neighbors. Using the nearest-neighbors approach, differential expression was determined by considering protein expression in a given cell type involved in a particular interaction, relative to cells not involved in those interactions. We first sought to understand the impact of Activated Tregs on protein expression in TOX+ CD8 T cells, given the enrichment of Activated Treg interactions with TCF1+ TOX+ CD8 T cells in the setting of recurrence (**Figure 5F - 5I**). TCF1- TOX+ CD8 T cells adjacent to Activated Tregs revealed decreased expression of granzyme B (GZMB) and interferon-γ (IFNG), with similar but nonsignificant pattern for TCF1+ TOX+ CD8 T cells adjacent to Activated Tregs (**Supplementary Figure 6A, 6B**). However, ICOS expression was not exclusive to Activated Tregs due to clustering and indeed many other Treg cells expressed ICOS in the SLN data (**Supplementary Figure 4D**). Thus, to better understand the influence of ICOS+ Tregs, we grouped all three Treg subsets together while retaining all other immune cell subsets.

We next used differential protein expression to understand the impact of neighboring Tregs on protein expression in CD8 T cells. Both TCF1- TOX+ and TCF1+ TOX+ CD8 T cells had lower expression of GZMB and IFNG when adjacent to a Treg compared to phenotypically matched CD8 T cells not adjacent to Tregs (**Figure 6A, 6B**). Conversely, expression of GZMB and IFNG was increased in TOX+ CD8 adjacent to Tumor cells, highlighting the contrast in protein expression depending on cellular neighbors (**Supplementary Figure 6C, 6D**). To assess protein expression at the per-cell level, we evaluated TOX+ CD8 T cells in the dataset relative to immediate proximity to Tregs. Indeed, GZMB and IFNG were reduced in TOX+ CD8 T cells adjacent to Treg irrespective of TCF1 expression (**Figure 6C, 6D**). Moreover, expression of GZMB and IFNG was assessed across all cell subsets within the SLN to understand whether Treg proximity was globally associated with reduced effector molecule expression. Most immune subsets, including GZMB CD8, did not show substantive changes in GZMB or IFNG expression based on proximity to Tregs. However, of the subsets that did have altered effector molecule expression given proximity to Treg, the most affected were TOX+ CD8 T cell subsets (**Figure 6E, 6F**). Next, we sought to determine whether the effect of Tregs on adjacent cells was dependent on Euclidean distance. To identify the nearest Treg for every cell, the spatial k-Nearest Neighbors (KNN) approach was broadened to capture a wider range of neighbors for each cell (k = 50) and the distance of the nearest Treg was determined as greater or less than 6 µm, representing a distance equivalent to one cell diameter (**Supplementary Figure 6E**). TCF1- TOX+ CD8 T cells within 6 µm of a Treg had lower expression of GZMB compared to TCF1- TOX+ CD8 T cells whose nearest Treg neighbor cells with more distant Treg neighbors, though this effect was less pronounced in the TCF1+ TOX+ CD8 T cell subset (**Figure 6G**). Conversely, both TCF1- and TCF1+ TOX+ CD8 T cells within 6 µm of a Treg neighbor exhibited higher expression of IFNG than TCF1- and TCF1+ TOX+ CD8 T cells with more distant Treg neighbors (**Figure 6H**). These data suggest that Treg adjacent to CD8 T cell subsets, particularly TOX+ subsets, are associated with altered expression of effector proteins, and that Treg associated reduction of GZMB expression within exhausted CD8 T cell subsets may predominantly occur over short distances.

**Figure 6.**
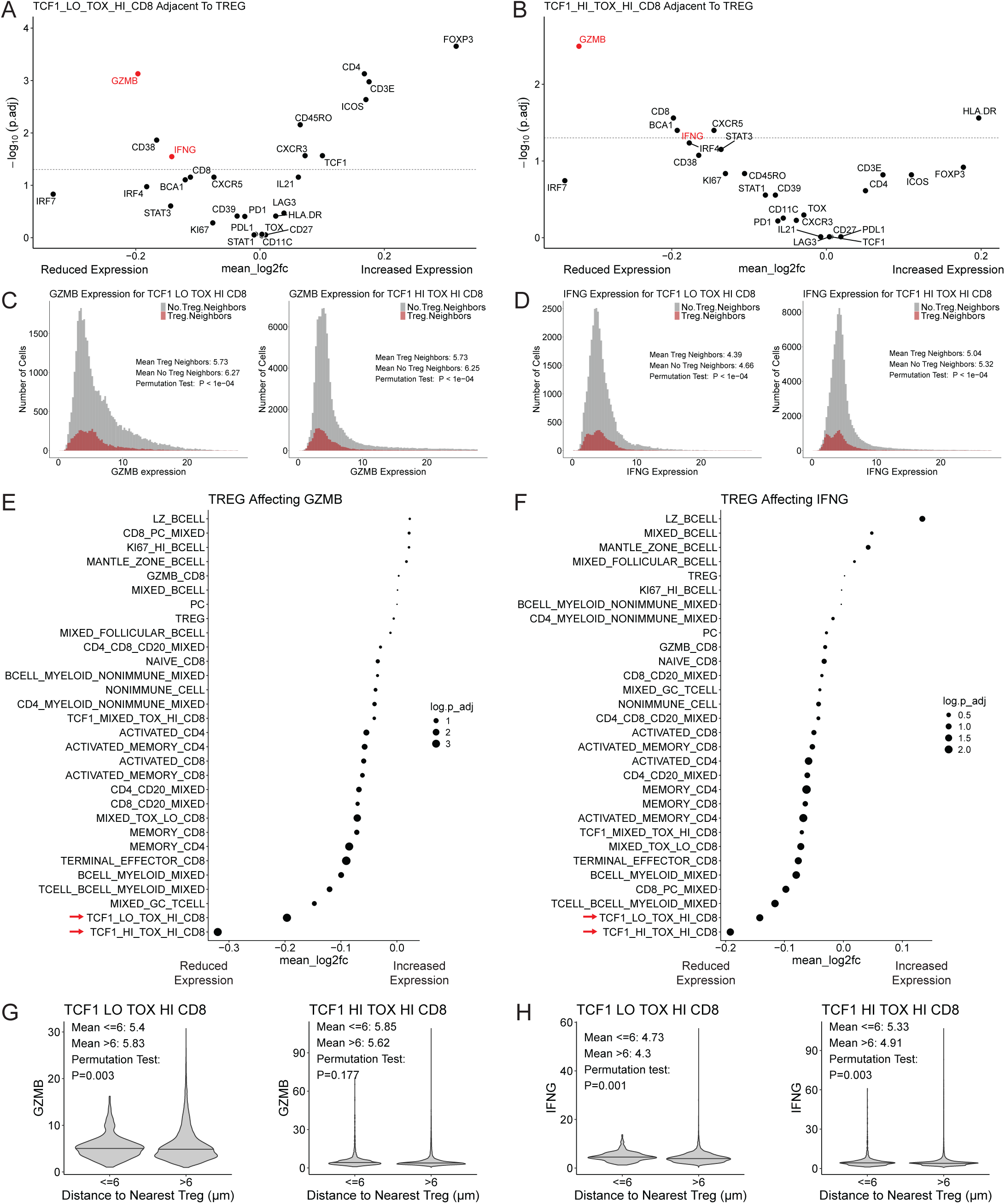
Proximal Tregs Influence Protein Expression of Exhausted CD8 T Cells in Melanoma SLN. **A-B**. Volcano plots displaying enrichment of protein expression in (A) TCF1 LO TOX HI CD8 or (B) TCF1 HI TOX HI CD8 adjacent to Tregs compared to subtype interactions with all other non-Treg cells. Dotted line at y=1.3 corresponding to P.adj=0.05. **C-D**. Histogram of (C) GZMB or (D) IFNG cell-level protein expression in TCF1 LO TOX HI CD8 and TCF1 HI TOX HI CD8 when adjacent to Treg neighbors (red) compared to adjacency to all other non-Treg cells (grey). **E-F**. Dot plot of (E) GZMB or (F) IFNG protein expression in cell subsets when found adjacent to Tregs compared to subtype interactions with all other non-Treg cells. **G-H**. Violin plots of (G) GZMB or (H) IFNG protein expression for TCF1 LO TOX HI CD8 and TCF1 HI TOX HI CD8 with Treg neighbors either > or ≤ 6 μm distance from index subtype. Y axis indicates z-score of expression.

Furthermore, Tregs undergoing activation demonstrate increased expression of proteins, such as LAG3, ICOS, and many others.^25,48–50^ Thus, given the decreased expression of effector molecules in TOX+ CD8 T cells neighboring Tregs, we sought to determine if Tregs involved within these interactions also had phenotypic alteration. Indeed, Tregs neighboring TCF1+ TOX+ CD8 T cells displayed increased expression of ICOS, PD-L1, FOXP3, LAG3, and CD39 compared to Tregs not involved in these pairwise interactions, though this effect was not observed for Tregs neighboring TCF1- TOX+ CD8 T cells (**Supplementary Figure 6F, 6G**). Whether functional inhibition of TCF1- TOX+ CD8 T cells occurs through distinct Treg molecular signaling pathways or from other modulatory signals within the SLN microenvironment will require further study. Nonetheless, these data establish a clear association between the physical proximity of Tregs and TOX+ exhausted CD8 T cells and suppression of CD8 T cell effector function, highlighting a spatially-regulated mechanism of immune modulation in the SLN.

## DISCUSSION

Despite great improvements in care of patients with melanoma over the past several decades, our understanding of the underlying mechanisms for T cell dysfunction remains incomplete. Sentinel lymph nodes (SLN) are routinely sampled to inform melanoma pathologic disease staging but are not routinely evaluated for state of immune cells and their interactions. Through high-dimensional spatial proteomic analysis, we characterized the immune landscape of clinical Stage I and Stage II melanoma SLN yielding an atlas of cellular states and interactions comprising nearly 12 million cells. Lymphoid region analysis revealed subtle shifts in regional cell abundance by stage, evidenced by higher frequencies of activated and memory CD4 T cells in the T Cell Dense and T/B Mixed regions of Stage I compared to Stage II SLN. Tissue-based transcriptional profiling implicated reduced T cell activity in Stage II SLN of patients who experienced disease recurrence. Thus we performed cellular interactivity profiling through a novel computational toolkit, Effect Size Interaction mapping (ESI-map), which demonstrated rewiring of pairwise cellular interactions, most notably involving ICOS-expressing Activated Tregs and TOX-expressing Exhausted CD8 T cells. Activated Tregs had enriched interactions with TCF1-expressing Exhausted CD8 T cells in Stage II patients who experience disease recurrence compared to those who remain recurrence free. Moreover, Exhausted CD8 T cells adjacent to Tregs had reduced functional activity, displaying decreased expression of effector molecules granzyme B (GZMB) and interferon-γ (IFNG). These findings build upon our understanding of how cellular spatial relationships in the SLN are related to anti-tumor immunity and the risk of melanoma clinical progression.

Spatial methodologies are being increasingly used to characterize cellular dynamics in the tumor microenvironment. Indeed, previous studies focusing on the melanoma primary tumor have similarly leveraged spatial techniques to identify prognostic biomarkers and decipher cellular relationships that drive anti-tumor immune responses. Interactions between tumor, stroma and immune cells defined by organization in cellular neighborhoods, colocalization in cellular compartments, and spatial density or distance of cellular phenotypes to each other have stratified melanoma patients by survival and response to immune checkpoint inhibitor (ICI) therapy.^32–36,51^ Furthermore, while limited studies have probed the spatial landscape of the melanoma tumor draining lymph nodes (TDLN), analyses of Stage III SLN revealed that interactions of tumor cells or cytotoxic lymphocytes with innate immune cells can predict melanoma recurrence outcomes.^52^ These studies provide insights on how immune cell interactions can inform patient outcomes, but additional insights into certain key lymphocyte subsets including exhausted CD8 T cells and Tregs are needed. Our study reveals that enriched interactions between Tregs and Exhausted CD8 T cells in the SLN may not only inform dynamics to anti-tumor immune responses, but also presents as a signature for melanoma disease recurrence. This spatial localization could be used to identify candidates for more intensive adjuvant ICI therapy by identifying patients with high risk of recurrence. These findings underscore the necessity for spatial approaches to fully understand the complexities of the anti-tumor immune response.

Beyond defining a prognostic signature based on cell-cell interactions, our study constructs a foundational spatial map of the SLN immune landscape across clinical Stage I and Stage II melanoma. This atlas is needed because SLN are frequently sampled for pathologic staging, but the immune environment of the SLN remains poorly understood. While previous studies have characterized the immune cell profile of dissociated SLN,^14,53^ dedicated spatially aware analyses are needed to resolve organizational patterns that are critical to immune cell function. Consistently across stage and outcome, several activated T cell populations were distributed within the T Cell Dense region, a critical site for antigen presentation and T cell survival. Additionally, small proportions of Activated Tregs and GZMB CD8 were found in and around the Tumor region in Stage II SLN. Tumor proximal Tregs may promote tumor growth and metastasis through local TGF-β signaling.^54–56^ In contrast, the association between proximity of cytotoxic CD8 T cells to melanoma cells and better ICI response suggests that a similar spatial biomarker could predict immunotherapy outcomes in the SLN.^35^ These findings provide novel insight into immune cell regional organization within the SLN, demonstrating that T cell positioning at key functional locations remains consistent despite advanced or recurrent disease.

The TDLN function as a reservoir for CD8 T cells, which are the precursors for intratumoral CD8 and are the key responders to ICI therapy.^5,15,17^ Biological processes that affect CD8 T cells may have important implications for success of ICI therapy. CD8 T cell inhibition due to Tregs may limit CD8 T cell migration and intratumoral responses as well as stunt patient response to ICI.^15,17,22,25,57^ Indeed, high density of Tregs in the SLN is associated with poor clinical response in melanoma.^45^ While we note proximity of Tregs to exhausted CD8 T cells affects CD8 T cell functional capacity, this relationship may be bidirectional as IL-2 production by CD8 T cells can enhance Treg immunomodulatory capacity by inducing ICOS expression.^58,59^ Unfortunately, the specific cellular interactions through which Tregs mediate immune modulation remain unclear, and systemic targeting of Tregs has been associated with variable benefit and sometimes substantial toxicity.^50,59^ Systematic evaluation of mechanisms for Treg suppression in the interactions with CD8 T cell subsets may identify specific proteomic or transcriptional pathways of interest, such as direct cell surface protein interactions or secretion of TGF-β.^50,60^ Directly targeting these pathways could limit unintended effects of Tregs and also enhance CD8 T cell efficacy.

Our data demonstrate a robust characterization of the melanoma SLN spatial immune landscape, though there are several opportunities for further insights. First, the scope of our analysis was inherently limited by the number and specificity of antibodies selected for our CODEX panel. We prioritized in-depth characterization of T and B lymphocytes, thus our study does not deeply interrogate innate or stromal cell populations which have been implicated in other melanoma studies.^32,33,61^ Second, some cells could not be resolved into distinct lineages likely due to technical challenges such as cell overlap within the tissue section and difficulty establishing precise membrane boundaries. This is a common limitation in multiplex immunofluorescence imaging that could be partially addressed with improved algorithms for cell segmentation and cell membrane spillover compensation, though physical overlap of cells in the tissue section cannot be easily resolved with bioinformatic approaches. Finally, while our dataset evaluated nearly 12 million cells, the ability to study patient-level factors such as tumor type, driver mutations, and demographics will require more patient samples to ensure adequate statistical power. Future studies in a larger patient cohort would help validate these findings and also allow for more robust evaluation of interactions involving multiple cell types.

Spatial proteomic characterization of Stage I and II melanoma SLN is a promising approach to improve understanding of how cell-cell interactions are altered with advanced disease and can be associated with clinical outcomes. Moreover, spatial immune profiling can provide opportunities for developing more carefully-targeted treatments and identifying new insights into disease mechanisms.

## METHODS

### Subject Details

FFPE SLN specimens from resectable melanoma patients were used for this study under IRB # 848619 in accordance with the institutional review board at the University of Pennsylvania.

### Antibody Screening

Antibodies for CODEX (Akoya Biosciences) were selected to profile a range of T and B lymphocytes and their signaling pathways, lymphoid structure, and tumor metastases based on literature review for the melanoma SLN. Criteria used to select antibodies of interest for screening include suitability for FFPE tissue and availability in anti-human, carrier-free, purified form. Each antibody was screened by indirect immunofluorescence (IF) on FFPE human tonsil tissue slides. Tonsil tissue was sectioned at 5 µm thickness, mounted onto charged slides and stored at room temperature. To melt the wax, slides were placed on a heating plate at 55 °C for 20 minutes. Following this, slides were deparaffinized in Histochoice clearing agent and rehydrated in descending concentrations of ethanol and ddH2O for 5 minutes each (Histochoice x 2, 100% EtOH x2, 90% EtOH, 70% EtOH, 50% EtOH, 30% EtOH, ddH2O x 2). Antigen retrieval was performed in 1X DAKO buffer (pH 9) with a pressure cooker for 20 minutes. Following this, samples were brought to room temperature and submerged in ddH2O x 4 (1st/3rd wash 5 seconds, 2nd/4th wash 1 minute). Slides were then washed 4 times in PBS for 5 minutes each. To prevent evaporation, a humidity chamber was assembled for slide incubations using an empty tip box filled with ddH2O and a paper towel submerged; slides are placed on top of the empty rack with the lid closed during incubations. Slides were blocked for 30 minutes inside the humidity chamber with 200 uL of a blocking solution containing 5% animal serum and 0.05% Tween 20 in PBS (PBS-T). After removing the blocking solution, slides were incubated for 2 hours with primary antibody diluted at a recommended concentration in 1% animal serum in PBS-T. Slides were then washed twice in PBS-T for 10 minutes each. Following this, slides were incubated for 1 hour with secondary antibodies diluted at 1:500 in 1% animal serum in PBS-T. Slides were then washed twice in PBS-T for 10 minutes each. A few drops of Prolong Gold Antifade Mountant with DAPI was applied to the tissue followed by a coverslip and slides were placed to dry in the dark overnight at room temperature (RT). After 24 hours, coverslips were sealed with CoverGrip Coverslip Sealant and imaged with the Keyence microscope used for subsequent CODEX imaging. Antibodies with appropriate cell-type expression and high signal-to-noise ratio based on image results were carried forward for conjugation to CODEX barcodes.

### Antibody Conjugation

A portion of the antibodies in the panel were purchased from Akoya in which they were pre-conjugated to CODEX barcodes. All other antibodies in the panel were custom conjugated to a CODEX barcode in our lab using the Akoya antibody conjugation kit in accordance to the Antibody Conjugation Protocol in the CODEX User Manual - Rev C. 50 kDA MWCO filters were blocked with 500 µL of filter blocking solution and centrifuged at 12000g for 2 minutes. After each step the liquid flow-through was discarded unless specified. Next 50 ug of antibody diluted in 100 uL of PBS was added to the filter and centrifuged at 12000g for 8 minutes. A reduction master mix was assembled with 6.6 µL of Reduction Solution 1 and 275 µL of Reduction Solution 2; 260 µL of reduction master mix was added to the filter, followed by gentle vortexing and a 30 minute incubation. After incubation the filter was centrifuged at 12000g for 8 minutes followed by addition of 450 µL of Conjugation Solution to the filter and centrifugation at 12000g for 8 minutes. The CODEX barcode was resuspended 10 µL of molecular grade water followed by 210 µL of conjugation solution. The full 220 µL barcode solution was added to the filter, gently vortexed and incubated for 2 hours. After incubation the filter was centrifuged at 12000g for 8 minutes. Next, 450 µL of Purification Solution was added to the filter followed by centrifugation at 12000g for 8 minutes. This step was repeated for a total of 3 purifications. Finally, 100 uL of antibody storage solution was added to the filter. The filter was inverted onto an empty microcentrifuge tube and centrifuged at 3000g for 2 minutes to collect the final antibody solution which was stored at 4 °C for downstream use.

### Coverslip Preparation

Coverslips were prepared at least 2 days prior to tissue sectioning in accordance to the FFPE Poly-L-Lysine Coverslip Preparation Protocol in the CODEX User Manual - Rev C. Coverslips were evenly distributed in a glass beaker and 0.1% poly-L-lysine solution was added to the beaker to fully cover the coverslips. The beaker was gently mixed and after ensuring no overlapping coverslips, it was then covered with parafilm and incubated overnight at RT. Following incubation, the poly-L-lysine was removed and ddH2O was added to the beaker and gently mixed for 30 seconds to wash the coverslips for a total of 7 washes. The coverslips were placed in a single layer on a paper towel to dry overnight before using for tissue sectioning.

### CODEX Staining Protocol

Staining of tissue for CODEX imaging was done using the Akoya antibody staining kit in accordance to the FFPE Tissue Staining Protocol in the CODEX User Manual - Rev C. FFPE Tissue samples were sectioned at 5 µm thickness and mounted onto Poly-L-Lysine coated coverslips (See Coverslip Preparation). Coverslips were placed on a heating plate at 55 °C for 20 minutes. Following this, coverslips were placed inside a coverglass staining rack for deparaffinization in Histochoice clearing agent and rehydration in descending concentrations of ethanol and ddH2O for 5 minutes each (Histochoice x 2, 100% EtOH x2, 90% EtOH, 70% EtOH, 50% EtOH, 30% EtOH, ddH2O x 2). Antigen retrieval was performed in 1X DAKO buffer (pH 9) with a pressure cooker for 20 minutes. Following this, samples were brought to room temperature and submerged in ddH2O x 4 (1st/3rd wash 5 seconds, 2nd/4th wash 1 minute). Coverslips were then washed 4 times in PBS for 5 minutes each. This was followed by 2 washes with Hydration Buffer for 2 minutes each and a 30 minute incubation with Staining Buffer. The antibody cocktail was prepared in a blocking buffer solution of 181 µL Staining Buffer, 4.75 µL of N Blocker, 4.75 µL of G Blocker, 4.75 µL of J Blocker and 4.75 µL of S Blocker per sample. Each antibody was added at the optimally determined dilution to the blocking solution such that the final volume for each sample was 200 µL. The coverslips were placed inside the humidity chamber and the antibody cocktail was applied to each coverslip, fully covering the tissue for a 3 hour incubation at RT. Following this, the coverslips were placed back into the coverglass staining rack and washed 2 times in Staining Buffer for 2 minutes each. Post-Staining Fixing Solution was prepared with a 1:10 dilution of 16% PFA and into Staining Buffer and coverslips were incubated in this solution for 10 minutes. Coverslips were then washed 3 times in PBS for 3 minutes each. Coverslips were incubated in Methanol on ice for 5 minutes, followed by 3 PBS washes for 5 seconds each. The final fixative solution was prepared by diluting 20 uL of Fixative Reagent into 1 mL of PBS. Coverslips were placed onto the humidity chamber and incubated with 200 µL of final fixative solution for 20 minutes. Finally, coverslips were washed 3 times in PBS for 5 seconds each before proceeding directly to the photobleaching protocol.

### CODEX Photobleaching Protocol

Immediately after completing the CODEX Staining Protocol, the CODEX Photobleaching Protocol was performed which is an adaptation from Du et al., 2019 and described in the Autofluorescence Quenching Protocol for CODEX.^40^ The photobleaching solution was prepared as a final working solution of 4.5% (w/v) hydrogen peroxide (H2O2) and 20 mM Sodium Hydroxide (NaOH) in PBS. Coverslips were placed in a single well in a 6-well plate and submerged in the photobleaching solution. The 6 well plate with the lid on was sandwiched in between two LED lamps and incubated for 45 minutes at RT. The coverslips were moved into a fresh well of photobleaching solution for a second incubation under the LED lamps for 45 minutes at RT. Following this the coverslips were washed 4 times in PBS for 5 minutes each and placed in Storage Buffer and stored at 4 °C in the dark until downstream imaging.

### CODEX Reporter Plate and Imaging

Before imaging, the CODEX reporter plate was prepared in accordance to the CODEX Reporter Plate Preparation Protocol in the CODEX User Manual - Rev C. Our 43-Plex Antibody Panel was arranged in an 18-cycle run including blank cycles at the first and last cycle (**Table S4**). A reporter stock solution was prepared for each coverslip with 244 µL of Nuclease free water, 30 µL of 10x CODEX Buffer, 25 µL of Assay Reagent, and 1 µL of Nuclear Stain. For each cycle, the reporter solution was prepared with 235 µL of stock solution and 5 µL of reporter. If a cycle had less than 3 reporters, 5 µL of reporter stock solution was supplemented to bring the total volume for each cycle to 250 µL. Blank cycles used 250 µL of reporter stock solution. The reporter solution for each cycle was transferred into the corresponding well on the 96-well plate which was then covered with a foil seal and stored at 4 °C until directly before starting the CODEX imaging set-up.

We utilized the CODEX Instrument system that integrated with the Keyence BZ-X810 microscope for imaging, following the BZ-X800 Image Acquisition Settings to set up each experimental run. A 20x/0.75 magnification was used to acquire the images along with the following fluorescent cubes: DAPI, EGFP (FITC/Cy2), CY3/TRITC, Cy5. Directly before imaging, buffers and reagents were prepared and transferred into the appropriate containers connected to the CODEX Instrument including Dimethyl Sulfoxide (DMSO), ddH2O, and 1X CODEX Buffer (10X CODEX Buffer diluted in ddH2O, filtered). The CODEX Run was set up in accordance with the CODEX Instrument Manager (CIM) prompts. A total of 6 Z-stacks were acquired with a pitch of 1.5 µm for each image. Single or Multi Regions of tissue were selected for each sample through the Stitching window with the region size varying for each sample as determined by the Set Center and Number of Images function with region size by X and Y tiles. After region selection, image acquisition began and could range from 20-72 hours per sample depending on region size and number. The resulting folders contained image tiles that were stitched together and processed for downstream application.

### CODEX Image Processing

After image acquisition of a CODEX experiment, the final QPTIFF files were generated using the CODEX Processor Version 1.8.3.14. Here, the raw images acquired from the CODEX Instrument system and Keyence Microscope are aligned and stitched across cycles and regions and corrected for inherent auto-fluorescence in the tissue. Here we selected Processing Options for Background Subtraction, Deconvolution, Extended Depth of Field, Shading Correction and Diagnostic Output. Depending on the region size and number, the image processing could take 24-48 hours per sample. After processing was completed, the final image combined all regions for an individual sample into a single QPTIFF with a XY resolution of 0.377 µm/pixel. This image was then visualized in QuPath and utilized for downstream analysis.^62^

### HALO Manual Artifact Exclusion

The .QPTIFF images generated by the CODEX experiments were first processed using HALO image analysis software (Version 3.6, Indica Labs). Images were first assessed for tissue areas with distorted or saturated signals that were not biologically relevant and are potentially caused by salt crystals present in the buffers delivered onto the tissue during imaging. Areas were marked for exclusion using the exclusion pen in the HALO annotation tools (**Supplemental Figure 1C**). These areas were then excluded from the downstream data export to prevent inaccurate performance of the segmentation algorithms or false interpretation of immunofluorescence signal.

### HALO AI Nuclear Segmentation

Cell segmentation was performed using the Nuclei Segmentation classifier algorithm and training with HALO AI (Version 3.6, Indica Labs). In accordance with the Using the Nuceli Seg (HALO AI) Network section of the HALO 3.5 Neural Network Classifiers User Guide, we first loaded the nuclear segmentation classifier Nuclei Seg (HALO AI) - FL v1.0.0. This algorithm was pretrained on different fluorescent images with DAPI staining and is used as a baseline for subsequent training from user defined annotations. We next added training regions from several tissue samples in which we used DAPI signal to define the boundaries of nuclei (DAPI+) and non-nuclei (DAPI-) using the pen tool in the HALO annotations tab (**Supplemental Figure 1D**). These annotations were then added to the appropriate classes in the Nuclei Seg Classifier for training, with DAPI+ annotations added to the class Nuclei and DAPI- annotations added to the class Background. A resolution of 0.25 µm/px and minimum object size of 0 µm^2^ was used for training. During training, performance of the classifier was assessed with the Real-Time tuning window on various areas of the tissue until satisfactory segmentation was achieved. The trained algorithm was saved and employed for nuclear segmentation in downstream analysis.

### HALO AI Lymphoid Region Segmentation

Our Lymphoid Region Segmentation algorithm was developed using the HALO AI DenseNet AI V2 classifier (Version 3.6, Indica Labs). In accordance to the Creating Tissue Mask Classifiers section of the HALO 3.5 Neural Network Classifiers User Guide, we first loaded the tissue mask classifier DenseNet V2 (HALO AI). We next defined each class as a lymphoid region of the SLN with the most general class defined first in the hierarchy and the most specific or nested class defined last in the hierarchy: Background (no tissue or limited cell density), T/B Mixed (high T and B cell density), B Cell Dense (high B cell density), T Cell Dense (high T cell density), Germinal Center (high density of CD21+, KI67+, PD-1+ cells), Tumor (MART1+ cells), Artifact (distorted or saturated signal). Following this we used antibody marker expression from all markers in our panel to guide boundary definition of training regions for each of our specified classes again using the pen tool in the HALO annotations tab (**Supplemental Figure 1E**).

Annotations were defined in several tumor and non-tumor containing samples. Additionally, boundaries of regions had deliberate overlap to ensure continuous tissue segmentation and to reaffirm hierarchy among regions. Regions positioned lower in the hierarchy were given priority in areas of overlap, for example Germinal Centers were identified within B Cell Dense regions. These annotations were then added to the appropriate classes in the DenseNet V2 Classifier for training. A resolution of 2.03 µm/px and minimum object size of 5.00 µm^2^ was used for training. During training, performance of the classifier was assessed with the Real-Time tuning (RTT) window on various areas of the tissue until satisfactory segmentation was achieved (**Supplemental Figure 1E**). The trained algorithm was saved and employed for lymphoid region segmentation in downstream analysis.

### HALO Antibody Expression Threshold with HighPlex Module

The antibody marker expression feature matrix was generated through the HALO HighPlex FL module (Version 4.2.5, Indica Labs). In accordance with the HALO 3.5 HighPlex FL Step-by-Step guide v4.2.3, we first loaded the HALO HighPlex FL module in the Analysis tab. We defined the Analysis Magnification as an Image Zoom of 1.5. We next used Autofill in the Dye Selection tab to allow HALO to load all 43 antibody markers into the module. In the Nuclear Detection tab we selected AI Custom for the Nuclear Segmentation Type and loaded our nuclear segmentation algorithm (See HALO AI Nuclear Segmentation). We used the following settings for nuclear detection: Minimum Nuclear Intensity of 0.095, Maximum Image Brightness of 1, Nuclear Segmentation Aggressiveness of 0.65, Nuclear Size of 8.3, 571.7, Minimum Nuclear Roundness of 0, Number of Nuclear Dyes of 1, Nuclear Dye 1 was DAPI-01 and Nuclear Dye 1 Weight of 1. In the Membrane and Cytoplasm Detection tab we set the Maximum Cytoplasm Radius to 1.5, Membrane Segmentation Aggressiveness to 0.97 and Cell Size to 10, 300. We did not define a Membrane Dye. For each marker, we individually defined either a Nuclear or Cytoplasm Positive Threshold and % Completeness Threshold based on satisfactory identification visualized within the Real-Time tuning window. This was completed independently for each sample in the cohort. In the Advanced tab we set Store Object (Cell) Data to True and set the Classifier Output Type to Mask. Here, we linked our Lymphoid Region Classifier to the Classifier Pipeline field (See HALO AI Lymphoid Region Segmentation). We then analyzed the image through the Analyze tab and selected Annotation Layer, ensuring that our annotation layer with the predefined exclusion areas were selected in the Annotations tab (See HALO Manual Artifact Exclusion). This generated interactive object data in the Results tab that included our single cell feature matrix with cell ID, antibody expression values, antibody threshold positive or negative classification (0 or 1), lymphoid region classification, and X,Y coordinates. We exported the object data as a .csv for each sample and utilized this data for downstream analysis.

### Nanostring

RNA was isolated from slides of FFPE SLN for analysis on the NanoString nCounter gene expression platform (NanoString Technologies). Before RNA isolation, tissue sections were deparaffinized in Xylene for 3 × 5 minutes, then sequentially rehydrated in 100% Ethanol for 2 × 2 minutes, 95% ethanol for 2 minutes, and 70% ethanol for 2 minutes, and then immersed in distilled H2O until ready to be processed. Tissue was lysed on the slide by adding 10–50 μL PKD buffer. Tissue was then scraped from the slide and transferred to a 1.5-mL Eppendorf tube. Proteinase K was added at no more than 10% final volume and the RNA lysate was incubated for 15 minutes at 55 °C and then 15 minutes at 80 °C. RNA was isolated using the Qiagen RNeasy FFPE kit per manufacturer protocol. Sample concentration was measured on the NanoDrop Spectrophotometer per manufacturer protocol. The resulting total RNA was stored at –80 °C until gene expression profiling was performed using the NanoString nCounter system. Per sample, 100 ng total RNA isolated from FFPE tissue was mixed with a 3′ biotinylated capture probe and a 5′ reporter probe tagged with a fluorescent barcode, from a custom-designed gene expression code set. Probes and lysate were hybridized overnight at 65 °C for 18 hours. Hybridized samples were then run on the NanoString preparation station using their high-sensitivity protocol per manufacturer instructions (NanoString Technologies). The samples were scanned at maximum scan resolution capabilities using the nCounter Digital Analyzer (NanoString Technologies).

All sample and data normalization occurred within the nCounter digital analyzer software, nSolver. Specifically, the raw code count data were normalized using a positive control normalization factor based on the spiked-in positive control raw counts and also a content normalization factor derived from the raw counts of a set of relevant housekeeping genes (ABCF1, C14ORF102, G6PD, OAZ1, POLR2A, SDHA, TBP, TBC1D10B, and UBB). Transcripts with counts less than or equal to the highest embedded negative controls (background noise) in that sample are first set to its background. The gene count for each gene is then subtracted from this background so that each sample has the same footing where zero numbers represent undetectable noise. Differential gene expression analysis was conducted with the limma package (3.2.1).^63^ Normalized counts were log2-transformed and a linear model was fitted for each gene followed by Empirical Bayes moderation. P-values were adjusted for multiple comparisons using the Benjamini-Hochberg method. Differentially expressed genes were determined by an adjusted p value of > 0.05 and a fold change of > 1.4. Gene set enrichment analysis was performed with Metascape.^64^

### Computational Analysis of Single-Cell Protein Expression Data

The single-cell protein feature matrix generated by HALO for each sample was used for downstream clustering analysis and visualization through the Giotto Suite (4.2.1).^41^ The feature matrices were cleaned by first extracting expression data for each marker as defined by Cell.Intensity and by generation of a single x,y coordinate for each cell by averaging the values for XMin/XMax and YMin/YMax as determined by HALO. For each sample a Giotto object was generated containing the metadata columns defined in the HALO output. Cells labeled as Background or Artifact by our Lymphoid Region Segmentation (Classifier.Label) were excluded from downstream analyses. The data were then scaled and normalized through normalizeGiotto with a standard normalization method (z-score). Following this, the Giotto objects were combined into a single Giotto object on a single x,y field through *joinGiottoObjects()*. To correct for sample variation, the combined Giotto object was further integrated though *runGiottoHarmony()*.^42^ Cells were clustered using *createNearestNetwork()* (dimensions_to_use = 1:10; k = 15) and *doLeidenClusterIgraph()* (resolution = 0.1; n_iterations = 100). Of the resulting 62 clusters, pan-negative clusters were identified as clusters below the 2nd percentile for mean total signal and excluded from downstream analyses. The remaining cells were again clustered using *createNearestNetwork()* (dimensions_to_use = 1:12; k = 15) and *doLeidenClusterIgraph()* (resolution = 0.1; n_iterations = 750) resulting in 64 unique clusters. Clusters with less than 100 cells were excluded from downstream analyses. The remaining cells were again clustered using *createNearestNetwork()* (dimensions_to_use = 1:12; k = 15) and *doLeidenClusterIgraph()* (resolution = 0.5; n_iterations = 500) resulting in 62 unique clusters. At this level, we classified clusters into the following groups: T Cells, B Cells, T/B Mixed Cells, Tumor or Artifact. Clusters labeled as Artifact were excluded from downstream analyses. Clusters labeled as Tumor were utilized in subsequent Combined Giotto Object and Neighborhood Analyses. All other T Cell, B Cell and T/B Mixed Cells were further characterized in our lymphoid cell subclustering strategy.

### Lymphoid Cell Characterization

The clusters identified as T Cells, B Cells and T/B Mixed Cells were used to generate 3 separate Giotto objects for downstream subclustering (**Supplemental Figure 2B**).

Harmony integration was run again on the B Cell Giotto Object which was then clustered using *createNearestNetwork()* (dimensions_to_use = 1:3; k = 15), *doLeidenClusterIgraph()* (resolution = 0.1; n_iterations = 200) and the following markers defined in clusterChannels: CD20, PAX5, HLA.DR, CD38, IGD, KI67, CXCR5, CD21, CD11C. The resulting 17 clusters within the B Cell Giotto Object were classified into the following groups based on marker expression: Mixed Follicular and Nonfollicular B Cell, Mixed Follicular B Cell, Nonimmune Cell, Plasma Cell (PC), Mantle Zone B Cell, KI67 High B Cell, B Cell/Myeloid/Nonimmune Cell Mixed.

Harmony integration was run again on the T/B Mixed Cell Giotto Object which was then clustered using *createNearestNetwork()* (dimensions_to_use = 1:5; k = 15), *doLeidenClusterIgraph()* (resolution = 0.3; n_iterations = 200) and the following markers defined in clusterChannels: CD20, CD19, PAX5, HLA.DR, CD38, IGD, KI67, CXCR5, CD21, CD11C, CD3E, CD4, CD8, TCF1, GZMB, CD39, PD1, TOX, FOXP3, ICOS, CD45RO. The resulting 26 clusters within the T/B Cell Giotto Object were classified into the following groups based on marker expression: Plasma Cell (PC), CD4/CD20 Mixed, CD8/CD20 Mixed, CD4/CD8/CD20 Mixed, CD8/PC Mixed, B Cell/Myeloid Mixed, T/B/Myeloid Mixed, Light Zone B Cell, Mixed GC T Cell, Artifact. Clusters labeled as Artifact were excluded from downstream analyses.

To separate out CD4 from CD8 T cells, cells in the T Cell Giotto Object with the HALO threshold classification of CD8 = 1 (See HALO Antibody Expression Threshold with HighPlex Module) were subclustered out as a CD8 T Cell Giotto Object. All other cells in the T Cell Giotto Object were subclustered out as a CD4 T Cell Giotto Object.

Harmony integration was run again on the CD8 T Cell Giotto Object which was then clustered using *createNearestNetwork()* (dimensions_to_use = 1:5; k = 15), *doLeidenClusterIgraph()* (resolution = 0.5; n_iterations = 200) and the following markers defined in clusterChannels: TCF1, GZMB, PD1, TOX, CD45RO, HLA.DR. In the resulting 36 clusters, 1 cluster was identified as Outlier and excluded from downstream analyses.

CD8 T Cells were further subclustered into 2 separate Giotto Objects based on either HALO threshold classification of TOX = 1 (TOX High) or TOX = 0 (TOX Low). Harmony integration was run again on both the CD8 TOX High and CD8 TOX Low Giotto Objects which were then individually clustered using *createNearestNetwork()* (dimensions_to_use = 1:3; k = 15), *doLeidenClusterIgraph()* (resolution = 0.1; n_iterations = 200) and the following markers defined in clusterChannels: TCF1, PD1, CD45RO, HLA.DR, GZMB. The CD8 TOX High Giotto Object resulted in 10 clusters that were classified into the following groups based on marker expression: TCF1 High TOX High CD8, TCF1 Low TOX High CD8, TCF1 Mixed TOX High CD8. The CD8 TOX Low Giotto Object resulted in 16 clusters that were classified into the following groups based on marker expression: Activated CD8, Activated Memory CD8, Granzyme B CD8, Naive CD8, Memory CD8, Terminal Effector CD8, Mixed TOX Low CD8.

Harmony integration was run again on the CD4 T Cell Giotto Object which was then clustered using *createNearestNetwork* (dimensions_to_use = 1:5; k = 15), doLeidenClusterIgraph (resolution = 0.5; n_iterations = 200) and the following markers defined in clusterChannels: CD3E, CD4, CD8, TCF1, GZMB, CD39, KI67, PD1, TOX, FOXP3, ICOS, CD45RO, HLA.DR. In the resulting 31 clusters, 1 cluster was identified as Outlier and excluded from downstream analyses.

CD4 T Cells were further subclustered into 2 separate Giotto Objects based on either HALO threshold classification of FOXP3 = 1 (FOXP3 High) or FOXP3 = 0 (FOXP3 Low). Harmony integration was run again on both the CD4 FOXP3 High and CD4 FOXP3 Low Giotto Objects which were then individually clustered using *createNearestNetwork()* (dimensions_to_use = 1:3; k = 15) and *doLeidenClusterIgraph()* (resolution = 0.1; n_iterations = 200). For clusterChannels, the CD4 FOXP3 High Giotto Object used FOXP3, CD39 and ICOS while the CD4 FOXP3 Low Giotto Object used TCF1, PD1, CD45RO, HLA.DR. The CD4 FOXP3 High Giotto Object resulted in 9 clusters that were classified into the following groups based on marker expression:

ICOS Low Treg, Activated Treg, Mixed Treg. The CD4 FOXP3 Low Giotto Object resulted in 16 clusters that were classified into the following groups based on marker expression: Activated CD4, Activated Memory CD4, Memory CD4, CD4/Myeloid/Nonimmune Cell Mixed.

### Cluster Identity Compilation in Main Giotto Object

Cluster annotations as well as any Artifact labels were extracted from the subclustered Giotto Objects (B Cell, T/B Mixed, CD8 TOX High, CD8 TOX Low, CD4 FOXP3 High, CD4 FOXP3 Low) and projected back onto the corresponding cells in the Combined Giotto Object. Clusters with identical names were combined. In total, we defined 33 unique clusters and excluded all cells labeled as Artifact or Outlier from our downstream analyses.

### Geojson Cluster and Region Projections in QuPath

To validate that our assigned cluster identities accurately align with the cellular phenotypes in the immunofluorescent images, we used the x,y coordinates in our dataset to generate GeoJSON files that project clusters as spatial annotations onto our CODEX images. For each sample, the x,y coordinates and the corresponding column with cluster labels were extracted into a matrix. For each unique value in the cluster column, st_multipoint from the sf (simple features) package was used to generate a single MultiPoint geometry object such that each cluster is represented as a collection of individual points based on x,y coordinates. We then utilized st_sf to generate a simple features object from the MultiPoint geometry object for each cluster. Each simple features object was saved as an individual GeoJSON file for each cluster. These files were then opened as annotations within the corresponding CODEX image in QuPath and used to visualize and validate cluster identities. Additionally, we employed this strategy to project annotations in QuPath for our Lymphoid Region Segmentation to visualize cellular phenotypes within specific lymphoid compartments.

### ESI-map Analyses

All spatial data processing and analysis were conducted in R (v4.4.3) using the Giotto suite (v4.2.2). Spatial Network was constructed to define cellular interactions. Two distinct methods were employed: a k-nearest neighbor (k-NN) approach and a fixed-radius approach. The primary k-NN network was built using the *createSpatialKNNnetwork()* function, connecting each cell to its 9 nearest neighbors (k=9). An alternative network was constructed to identify all neighbors within a 6 µm radius by setting a high k (k=50) and maximum_distance to 15.92. To determine if observed cellular interactions occurred more frequently than expected by chance, a permutation-based enrichment analysis was performed using the *cellProximityEnrichment()* function in Giotto suite. For each pairwise cell-type interaction, the observed frequency within the k-NN (k=9) network was compared against a null distribution generated by 200 random permutations of the cell-type labels. A log2 enrichment score was calculated as the log2 ratio of the observed interaction count plus a pseudocount of 1 to the average permuted interaction count plus a pseudocount of 1. P-values were calculated based on the permutation results and adjusted for multiple comparisons using the False Discovery Rate (FDR) method. These analyses were performed on a per-sample basis and subsequently aggregated by clinical metadata (e.g., disease stage, recurrence status).

The local cellular neighborhood, or niche, for each cell was defined as the proportional representation of its neighboring cell types within the k-NN (k=9) network. For specific cell types of interest, the average niche composition was calculated by averaging these proportions across all cells of that type within a sample or across samples within a clinical subgroup. Differences in niche composition between clinical conditions (Stage I vs. Stage II; Recurrence vs. No Recurrence) were visualized using dot plots and network diagrams generated with the ggplot2 (v3.5.2) and ggraph (v2.2.1) packages.

To investigate the influence of cellular location on gene expression, the distance from specific CD8+ T cell subsets to the nearest regulatory T cell (Treg) was calculated. A spatial network of all cells within each sample was constructed using a high k (k=50) to ensure all potential connections were captured. From this network, the shortest Euclidean distance from each TCF1_LO_TOX_HI_CD8 T cell to any Treg was identified. These cells were binned based on this distance (e.g., ≤ 6 µm vs. > 6 µm). Normalized expression values for features of interest, such as GZMB and IFNG, were retrieved from the Giotto object for these binned cell populations. The statistical significance of expression differences between distance bins was assessed using a permutation test with 10,000 iterations. To identify proteins whose expression is altered by cell-cell interactions,we performed the *findInteractionChangedFeats()* function in Giotto to compare protein features for each pairwise cell-type interaction defined by the k-NN (k=9) spatial network. A permutation test was used to assess the statistical significance of expression changes, and the resulting p-values were FDR-adjusted. This yielded a log2 fold change (log2FC) for each feature. For a consolidated analysis, all CD4 regulatory T cell (Treg) subtypes were grouped into a single ’TREG’ annotation. All analyses were performed using R packages including data.table (v1.17.6), dplyr (v1.1.4), and rstatix (v0.7.2) for statistical comparisons.

### Statistical Analyses

Permutation tests were used to compare distributions where indicated. All tests were performed as two-tailed tests intended to constrain Type I error at 0.05. Effect size was assessed as the difference between group means divided by the pooled standard deviation.

## TABLES

**Table S1.** Participant Demographics Full Cohort.

|  | **Overall** | **Stage I** | **Stage II SLN-** | **Stage II SLN+** |
| --- | --- | --- | --- | --- |
| **Number of participants** | 51 | 8 | 22 | 21 |
| **Age** |  |  |  |  |
| Median | 61 | 65 | 62 | 57 |
| **Sex (% Female)** | 24% | 63% | 23% | 9.5% |
| **Recurrence (% No)** | 57% | 100% | 59% | 38% |
| **Breslows Depth (mm)** | 3.7 | 0.7 | 4.5 | 3.7 |
| **Mitosis (mm<sup>2</sup>)** | 6 | 1 | 7 | 9 |
| **Ulceration (% Yes)** | 69% | 0% | 68% | 95% |
| **Tumor Size (mm<sup>2</sup>)** | 0.08 | NA | NA | 0.08 |
| Unknown | 38 | 8 | 22 | 8 |

**Table S2.** Participant Demographics Cohort for CODEX.

|  | **Overall** | **Stage I** | **Stage II SLN+** |
| --- | --- | --- | --- |
| **Number of participants** | 15 | 5 | 10 |
| **Age** |  |  |  |
| Median | 64 | 64 | 63 |
| Range | 36 – 74 | 50 – 68 | 36 – 74 |
| **Sex (% Female)** | 40% | 60% | 30% |
| **Recurrence (% No)** | 60% | 100% | 40% |
| **Breslows Depth (mm)** | 3.2 | 0.60 | 4.4 |
| **Mitosis (mm<sup>2</sup>)** | 8 | 1 | 12 |
| **Ulceration (% Yes)** | 70% | 0% | 70% |
| **Tumor Size (mm<sup>2</sup>)** | 0.37 | NA | 0.37 |
| Unknown | 10 | 5 | 5 |

**Table S3.** CODEX Antibody Panel.

| Antibody | Clone | Dilution | Barcode | Fluorescence Channel | Akoya or Custom Conjugation | General Cellular Phenotype |
| --- | --- | --- | --- | --- | --- | --- |
| CD20 | L26 | 1:200 | BX007 | AF488 | Akoya | B Cell |
| PAX5 | 1H9 | 1:100 | BX040 | AF488 | Custom | B Cell |
| Ki67 | B56 | 1:200 | BX016 | AF488 | Custom | T/B Cell |
| STAT3 | 124H6 | 1:200 | BX001 | AF488 | Custom | Signaling |
| IRF4 | IRF4.3E4 | 1:50 | BX004 | AF488 | Custom | B Cell |
| Vimentin | RV202 | 1:400 | BX010 | AF488 | Custom | Non-Immune Cell |
| Pan-cytokeratin | AE-1/AE-3 | 1:1000 | BX019 | AF488 | Akoya | Non-Immune Cell |
| Mac2/Gal3 | M3/38 | 1:400 | BX025 | AF488 | Custom | Non-Immune Cell/Tumor |
| S100B | E7C3A | 1:50 | BX037 | AF488 | Custom | Non-Immune Cell/Tumor |
| IRF7 | 12G9A36 | 1:50 | BX049 | AF488 | Custom | B Cell |
| STAT1 | D1K9Y | 1:200 | BX034 | AF488 | Custom | Signaling |
| SOX10 | D5V9L | 1:50 | BX031 | AF488 | Custom | Tumor |
| CXCR5 | EPR23463-30 | 1:50 | BX035 | ATTO 550 | Custom | T/B Cell |
| PMTOR | 49F9 | 1:50 | BX005 | ATTO 550 | Custom | Signaling |
| BCA1 | EPR23400-92 | 1:50 | BX002 | ATTO 550 | Custom | T/B Cell |
| TBET | D6N8B | 1:200 | BX052 | ATTO 550 | Akoya | T Cell |
| CD45RO | UCHL1 | 1:200 | BX017 | ATTO 550 | Akoya | T Cell |
| CD38 | E7Z8C | 1:800 | BX089 | ATTO 550 | Akoya | B Cell |
| CD21 | EP3093 | 1:2000 | BX032 | ATTO 550 | Akoya | B Cell/FDC |
| CD8 | CB/144B | 1:800 | BX026 | ATTO 550 | Akoya | T Cell |
| GZMB | D6E9W | 1:200 | BX041 | ATTO 550 | Akoya | T Cell |
| IFNG | EPR21704 | 1:400 | BX020 | ATTO 550 | Akoya | T Cell |
| TOX | E6I3Q | 1:200 | BX060 | ATTO 550 | Akoya | T Cell |
| CD45 | D9M81 | 1:200 | BX021 | ATTO 550 | Akoya | T/B Cell |
| CD39 | EPR20627 | 1:200 | BX099 | ATTO 550 | Akoya | T Cell |
| IL21R | Polyclonal | 1:50 | BX014 | ATTO 550 | Custom | Signaling |
| CD27 | M-T271 | 1:50 | BX023 | ATTO 550 | Custom | B Cell |
| IgD | W18340A | 1:50 | BX028 | ATTO 550 | Custom | B Cell |
| IL21 | 14k5H3 | 1:50 | BX050 | AF647 | Custom | Signaling |
| LAG3 | D2G40 | 1:50 | BX006 | AF647 | Custom | T Cell |
| FOXP3 | 236A/E7 | 1:50 | BX036 | AF647 | Custom | T Cell |
| CD4 | EPR6855 | 1:200 | BX003 | AF647 | Akoya | T Cell |
| CD3e | EP449E | 1:200 | BX045 | AF647 | Akoya | T Cell |
| PD1 | D4W2J | 1:200 | BX046 | AF647 | Akoya | T Cell |
| ICOS | D1K2T | 1:200 | BX054 | AF647 | Akoya | T Cell |
| PDL1 | 73-10 | 1:200 | BX043 | AF647 | Akoya | B Cell |
| Tcf1 | C63D9 | 1:200 | BX061 | AF647 | Akoya | T Cell |
| CD19 | LE-CD19 | 1:200 | BX042 | AF647 | Custom | B Cell |
| CD11c | 118/A5 | 1:200 | BX024 | AF647 | Akoya | B Cell |
| HLA-DR | EPR3692 | 1:1000 | BX033 | AF647 | Akoya | T/B Cell |
| CXCR3 | G025H7 | 1:800 | BX030 | AF647 | Custom | T Cell |
| GP100 | HMB45 | 1:200 | BX094 | AF647 | Akoya | Tumor |
| MART1 | EPR20380 | 1:200 | BX055 | AF647 | Custom | Tumor |

**Table S4.**
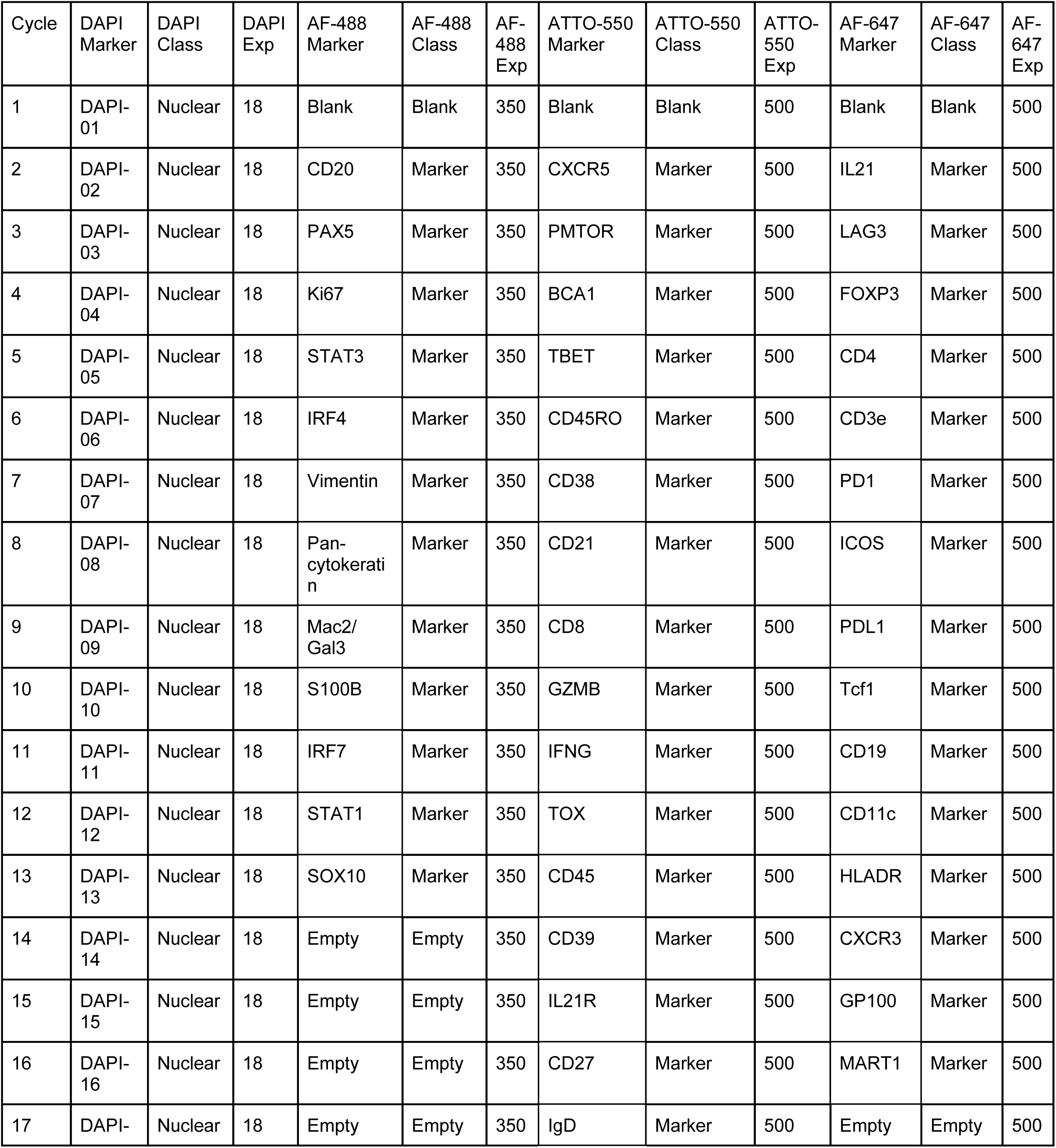
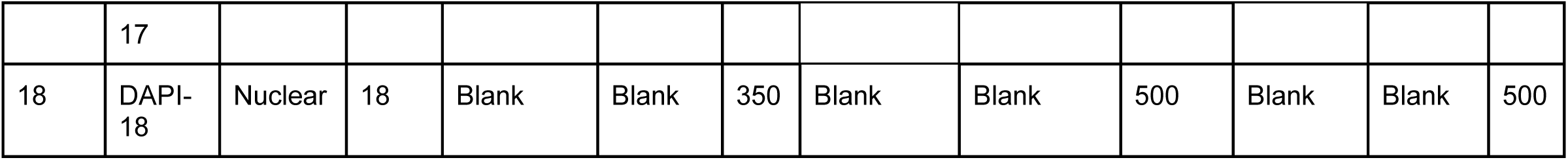
CODEX Experiment Set Up.

**Table S5.** CODEX Image Area and Cells Acquired.

| Sample | Stage | Total Area of Analyzed Tissue (mm<sup>2</sup>) | Total Cell Count Before Artifact Exclusion | Total Cell Count After Artifact Exclusion |
| --- | --- | --- | --- | --- |
| 301 | II | 9.743 | 202289 | 191806 |
| 302 | II | 50.02285 | 1130154 | 1085310 |
| 303 | II | 60.69677 | 1194595 | 1092977 |
| 304 | II | 5.91026 | 118574 | 110921 |
| 305 | II | 15.4114 | 293172 | 268034 |
| 306 | II | 100.14814 | 2051720 | 1992367 |
| 307 | II | 32.28168 | 701426 | 684647 |
| 308 | II | 45.74794 | 1069814 | 1042704 |
| 309 | II | 45.2687 | 959316 | 940962 |
| 310 | II | 49.98302 | 957919 | 936951 |
| 101 | I | 27.17739 | 537811 | 506512 |
| 102 | I | 37.47169 | 778998 | 743501 |
| 103 | I | 44.7109 | 942538 | 924650 |
| 104 | I | 30.90714 | 657623 | 640375 |
| 105 | I | 27.41166 | 575204 | 531238 |
| All Samples |  |  | 12171153 | 11692955 |

**Table S6.** Cell Type Annotation Definitions.

| Cluster | Protein Expression | Defining Spatial Features | Other |
| --- | --- | --- | --- |
| ACTIVATED TREG | CD4+, FOXP3+, ICOS+ |  |  |
| ICOS LO TREG | CD4+, FOXP3+, ICOS LO/- |  |  |
| MIXED TREG | CD4+, FOXP3+, ICOS +/- |  |  |
| MEMORY CD4 | CD4+, CD45RO+ |  |  |
| ACTIVATED CD4 | CD4+, HLADR+ |  |  |
| ACTIVATED MEMORY CD4 | CD4+, CD45RO+, HLADR+ |  |  |
| ACTIVATED CD8 | CD8+, HLADR+ |  |  |
| ACTIVATED MEMORY CD8 | CD8+, HLADR+, CD45RO+ |  |  |
| MEMORY CD8 | CD8+, CD45RO+ |  |  |
| GZMB CD8 | CD8+, GZMB+ |  |  |
| TERMINAL EFFECTOR CD8 | CD8+, TCF1 LO/-, CD45RO LO/-, CD3E LO |  |  |
| MIXED TOX LO CD8 | CD8+, TOX LO/-, CD45RO +/-, TCF1 +/- |  |  |
| NAIVE CD8 | CD8+, CD45RO-, TCF1+ |  |  |
| TCF1 LO TOX HI CD8 | CD8+, TOX+, TCF1 LO/- |  |  |
| TCF1 HI TOX HI CD8 | CD8+, TOX+, TCF1+ |  |  |
| TCF1 MIXED TOX HI CD8 | CD8+, TOX+, TCF1 +/- |  |  |
| MANTLE ZONE BCELL | CD20+, IGD+, CXCR5+ | Peripheral to GC |  |
| LZ BCELL | CD20+, KI67 LO/- | Polarized within GC |  |
| KI67 HI BCELL | CD20+, KI67+ |  |  |
| MIXED FOLLICULAR BCELL | CD20+ | Localized to B Cell Follicle |  |
| MIXED BCELL | CD20+ |  |  |
| PC (PLASMA CELL) | CD20-, CD3E-, CD38+, IRF4+ |  |  |
| CD8 PC MIXED | CD8+ AND/OR CD20-, CD3E-, CD38+, IRF4+ |  |  |
| CD8 CD20 MIXED | CD8+ AND/OR CD20+ |  |  |
| CD4 CD8 CD20 MIXED | CD8+ AND/OR CD4+ AND/OR CD20+ |  |  |
| CD4 CD20 MIXED | CD4+ AND/OR CD20+ |  |  |
| MIXED GC TCELL | CD3E+, CD4+ AND/OR CD8+, PD1 +/-, FOXP3 +/- | Localized to GC | Contained Tfh and Tregs |
| CD4 MYELOID<br>NONIMMUNE MIXED | CD4+ AND/OR MAC2/GAL3+ |  | Contained some Tumor cells |
| TCELL BCELL MYELOID<br>MIXED | CD3E+ AND/OR CD20+ AND/OR CD11C+<br>AND/OR MAC2/GAL3+ |  |  |
| BCELL MYELOID MIXED | CD20+ AND/OR CD11C+ |  |  |
| BCELL MYELOID<br>NONIMMUNE MIXED | CD20+ AND/OR MAC2/GAL3+ |  | Contained some Tumor cells |
| NONIMMUNE CELL | CD45-, CD20-, CD3E-, CD38-, MAC2/GAL3 +/- |  | Contained some Tumor cells |
| Tumor | CD45-, MART1+, GP100+/-, S100B +/-,<br>MAC2/GAL3 +/- |  |  |

## METHODS / KEY RESOURCES TABLE

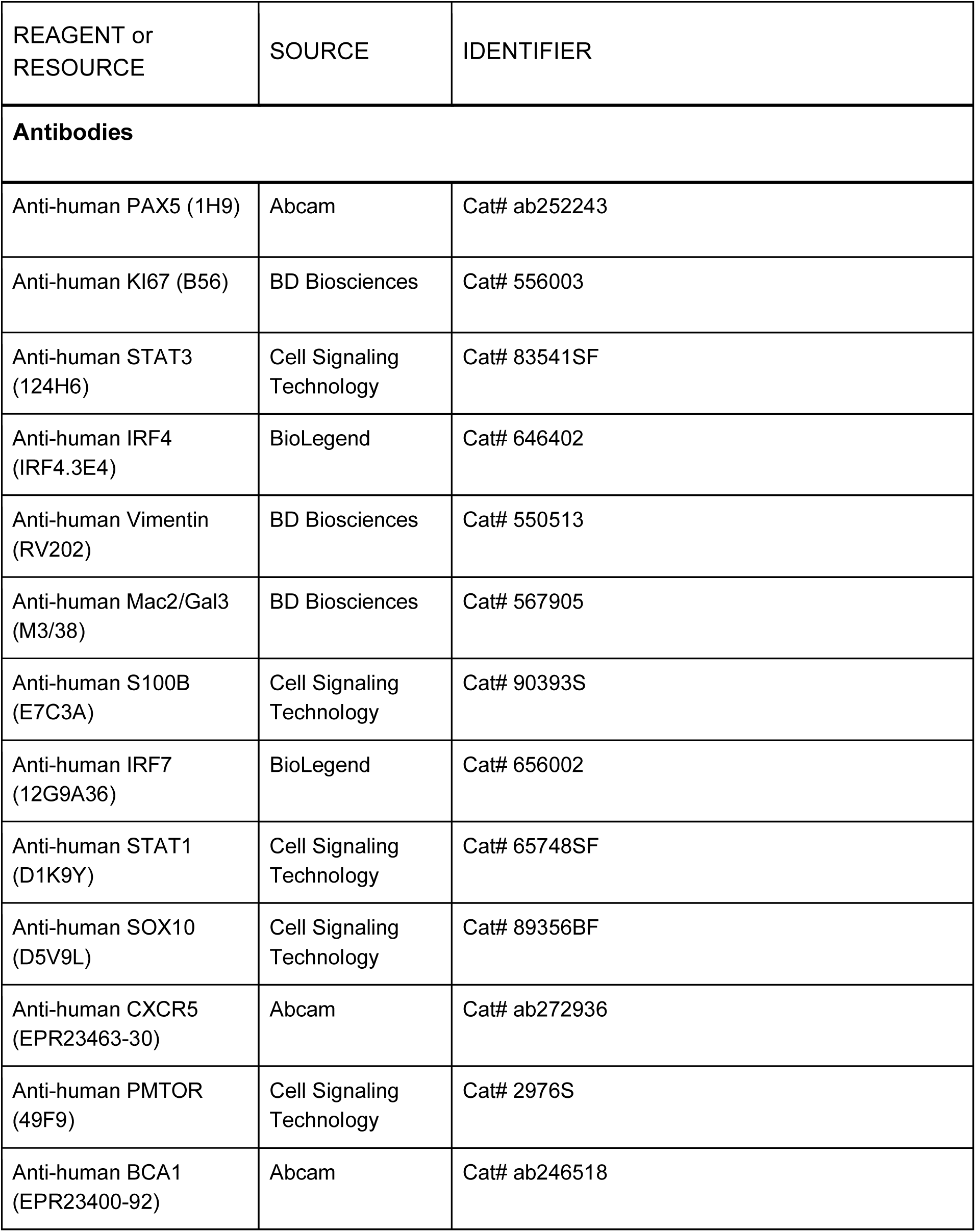
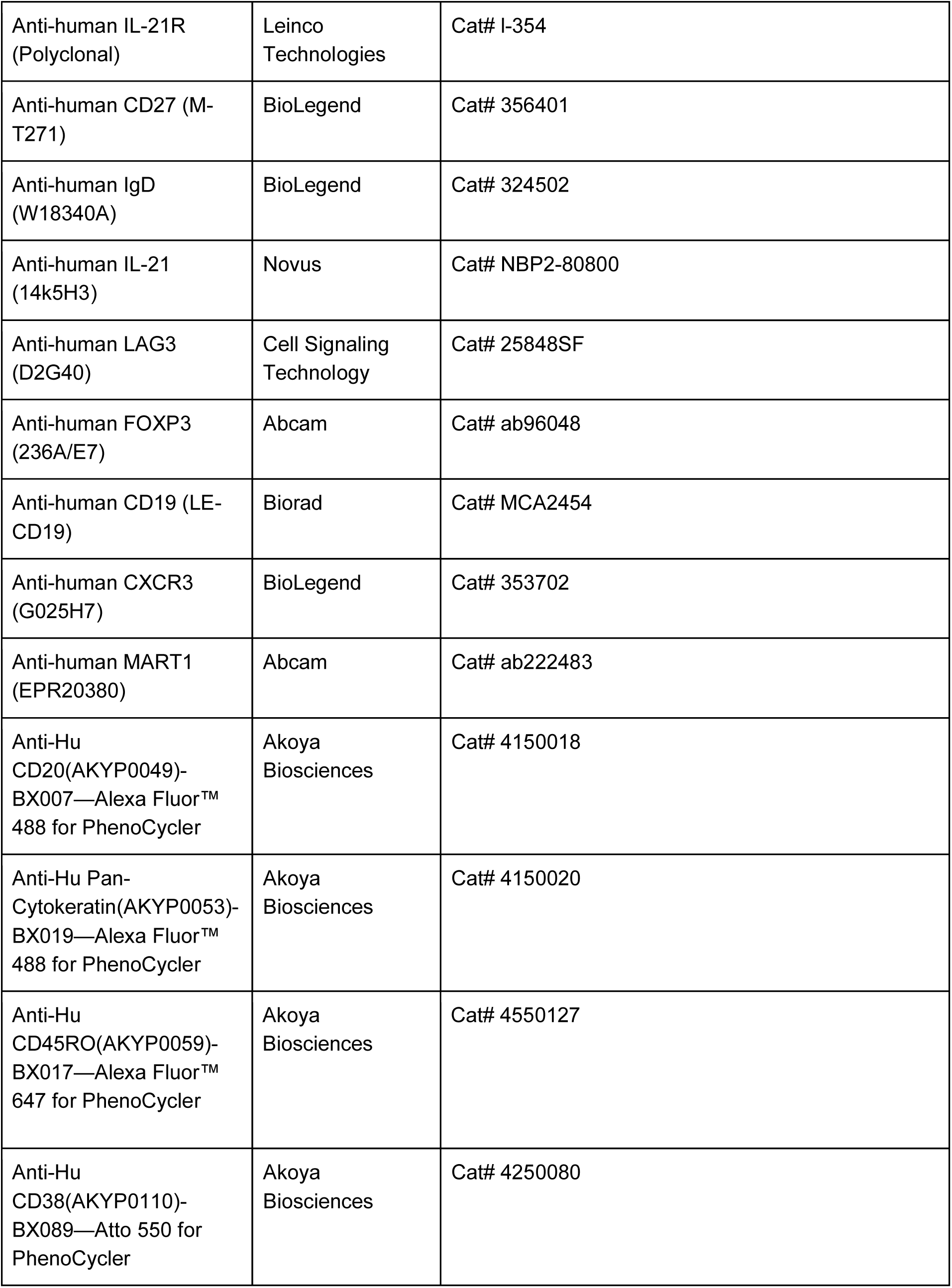
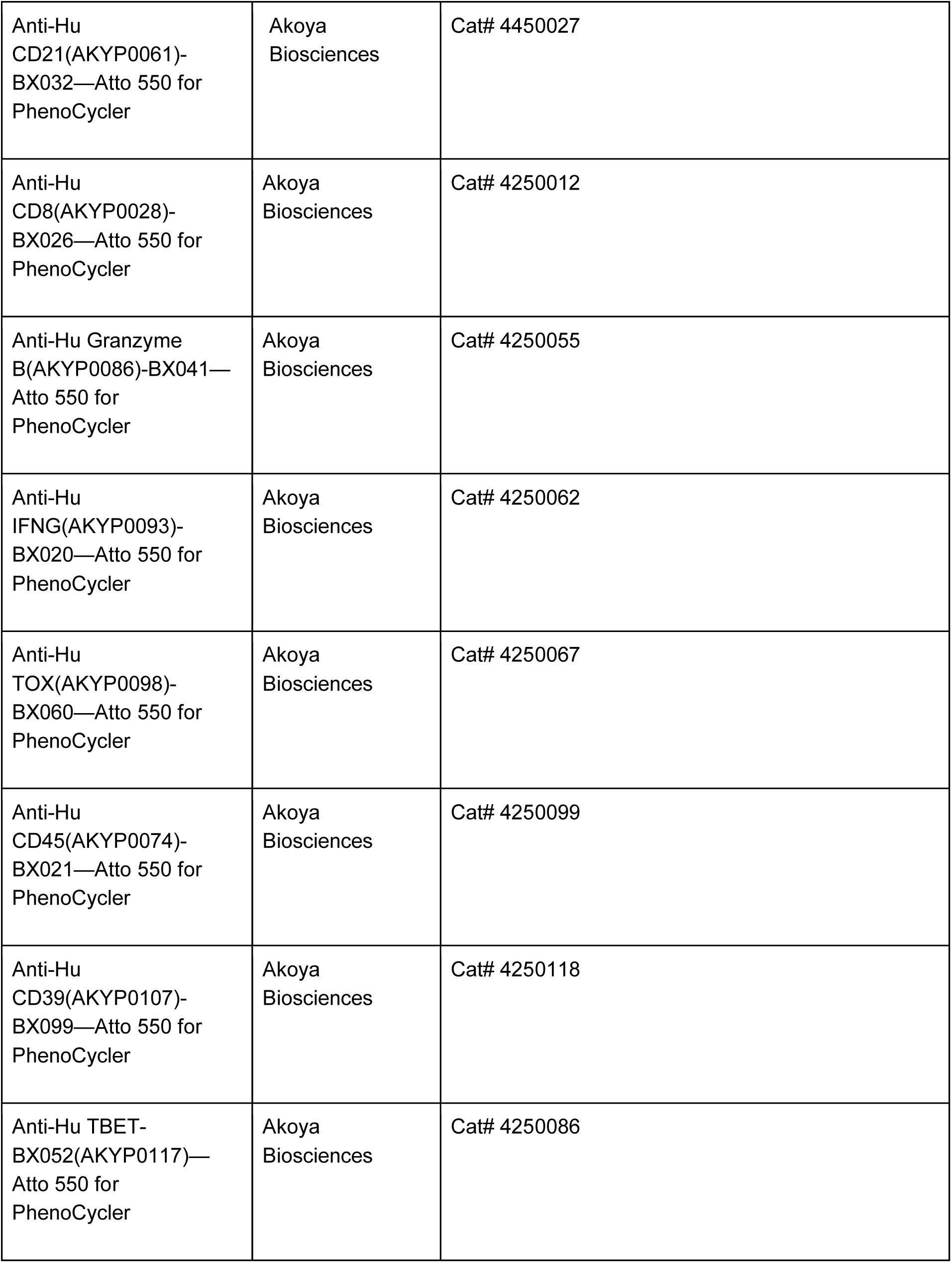
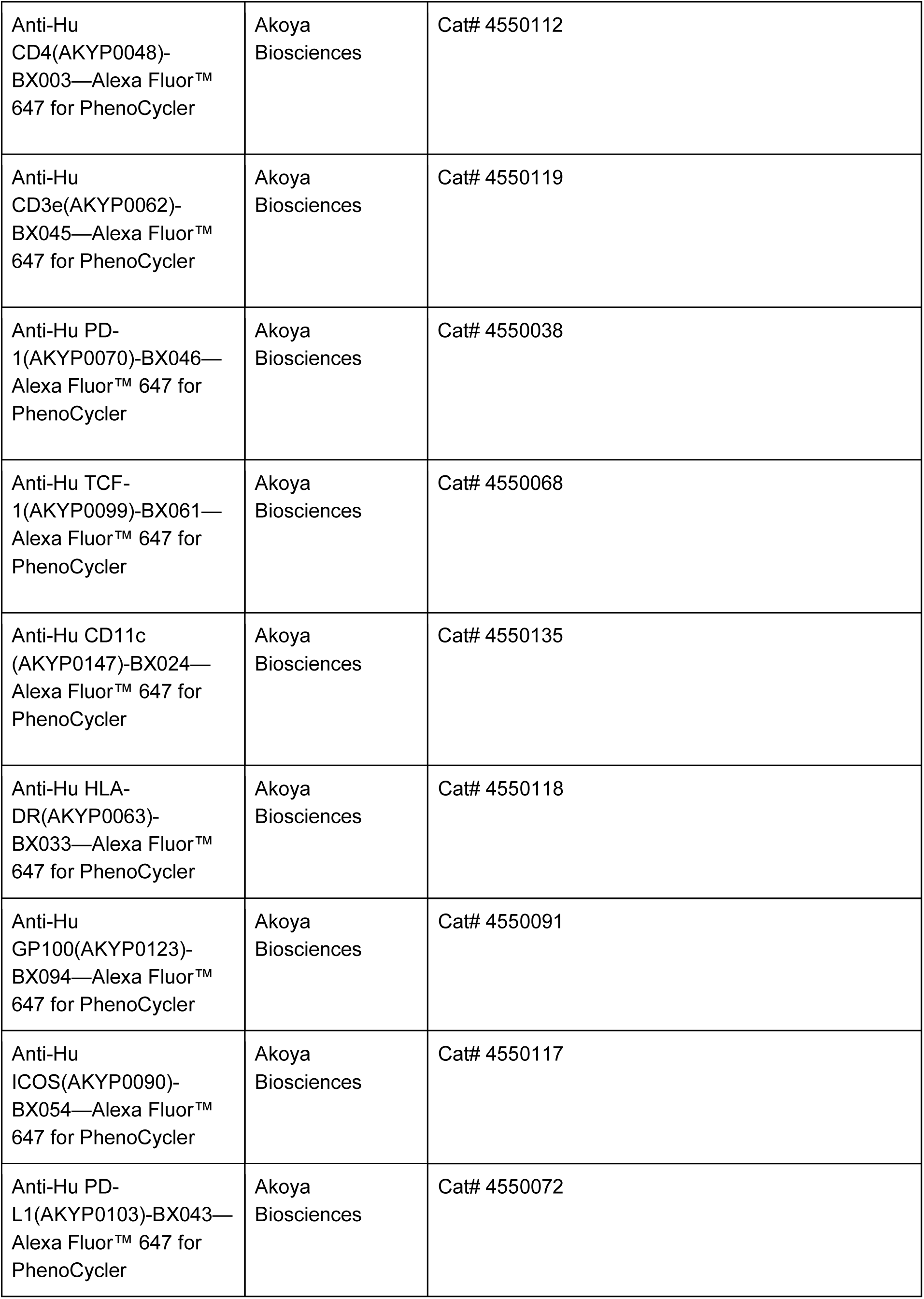
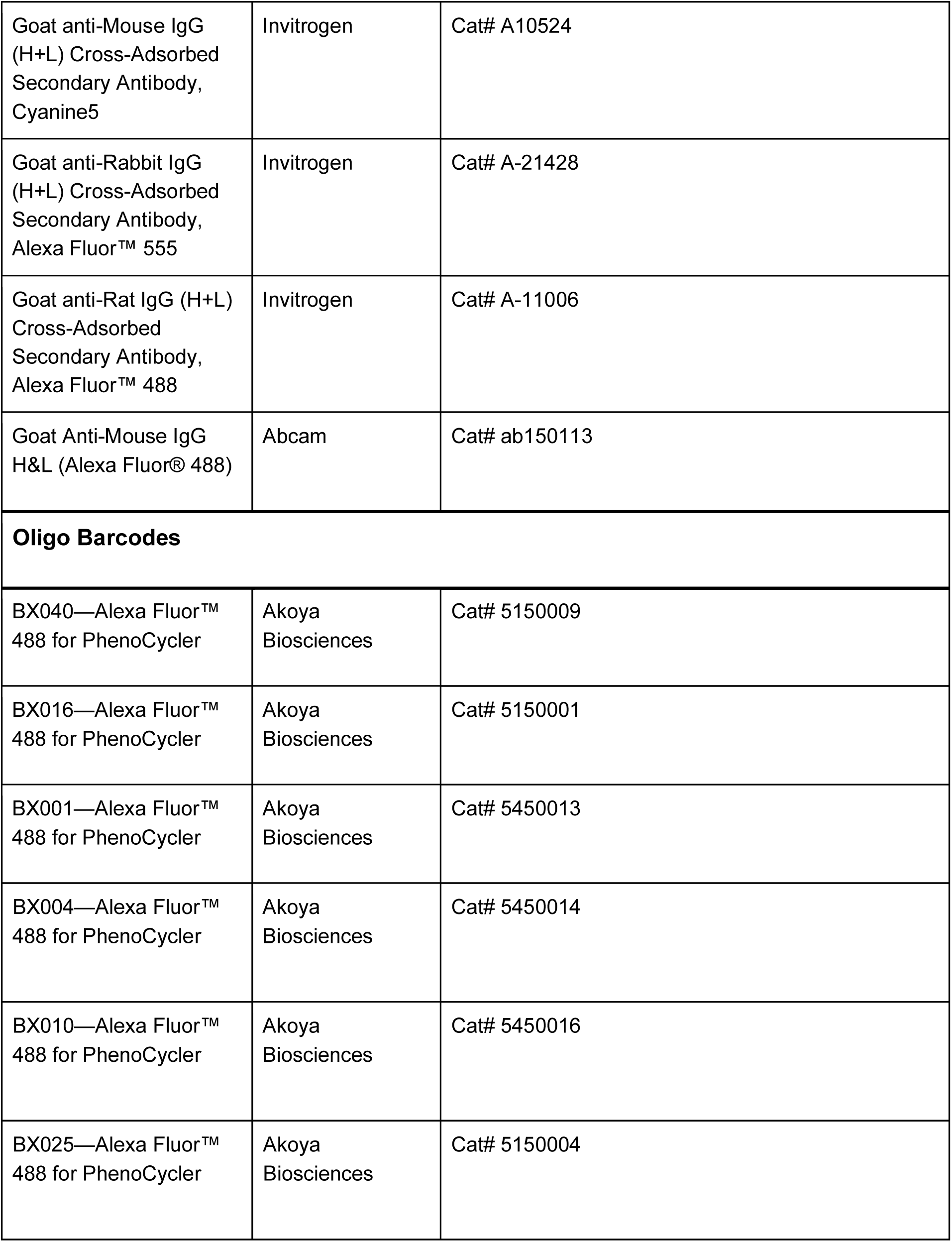
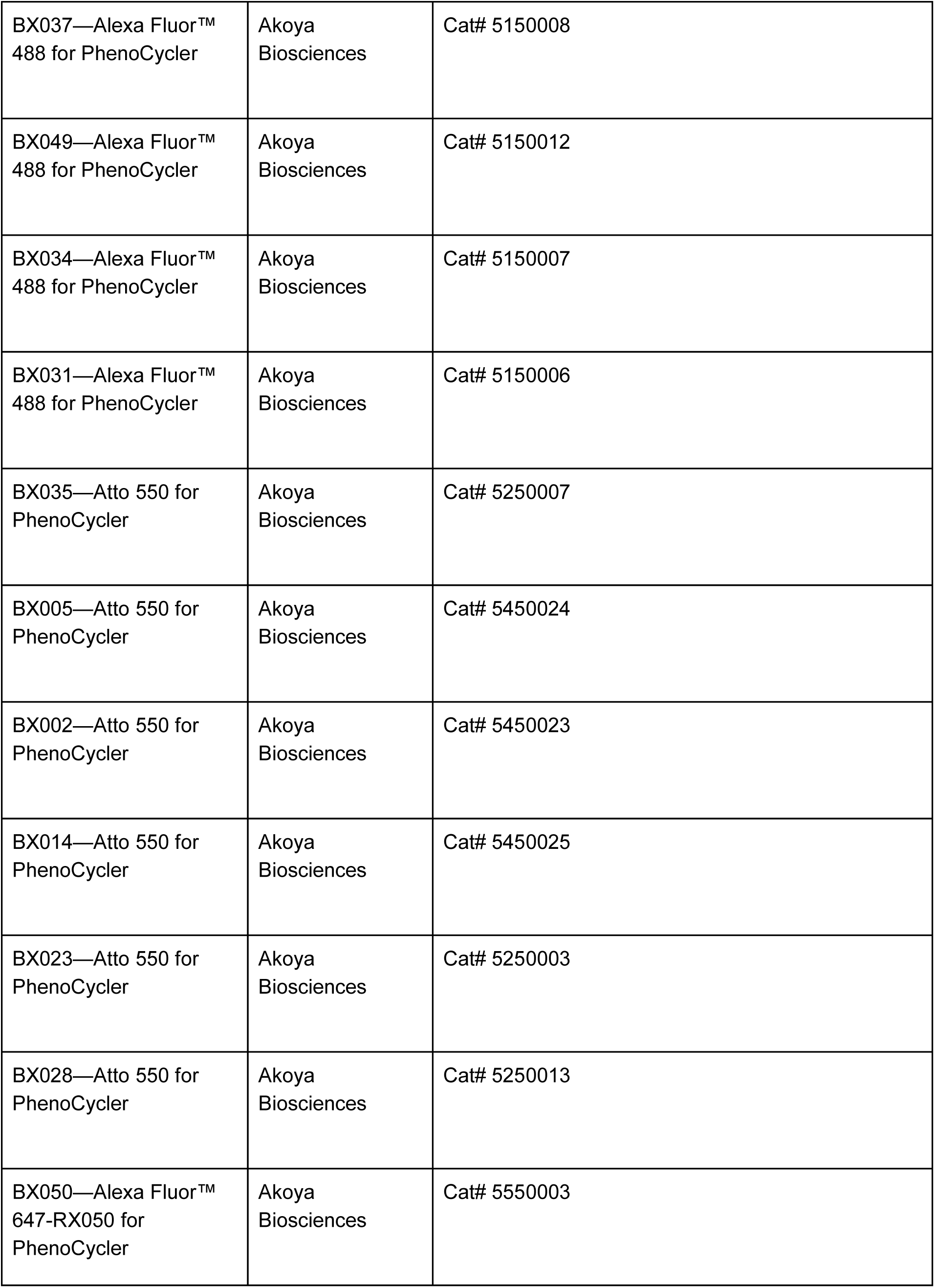
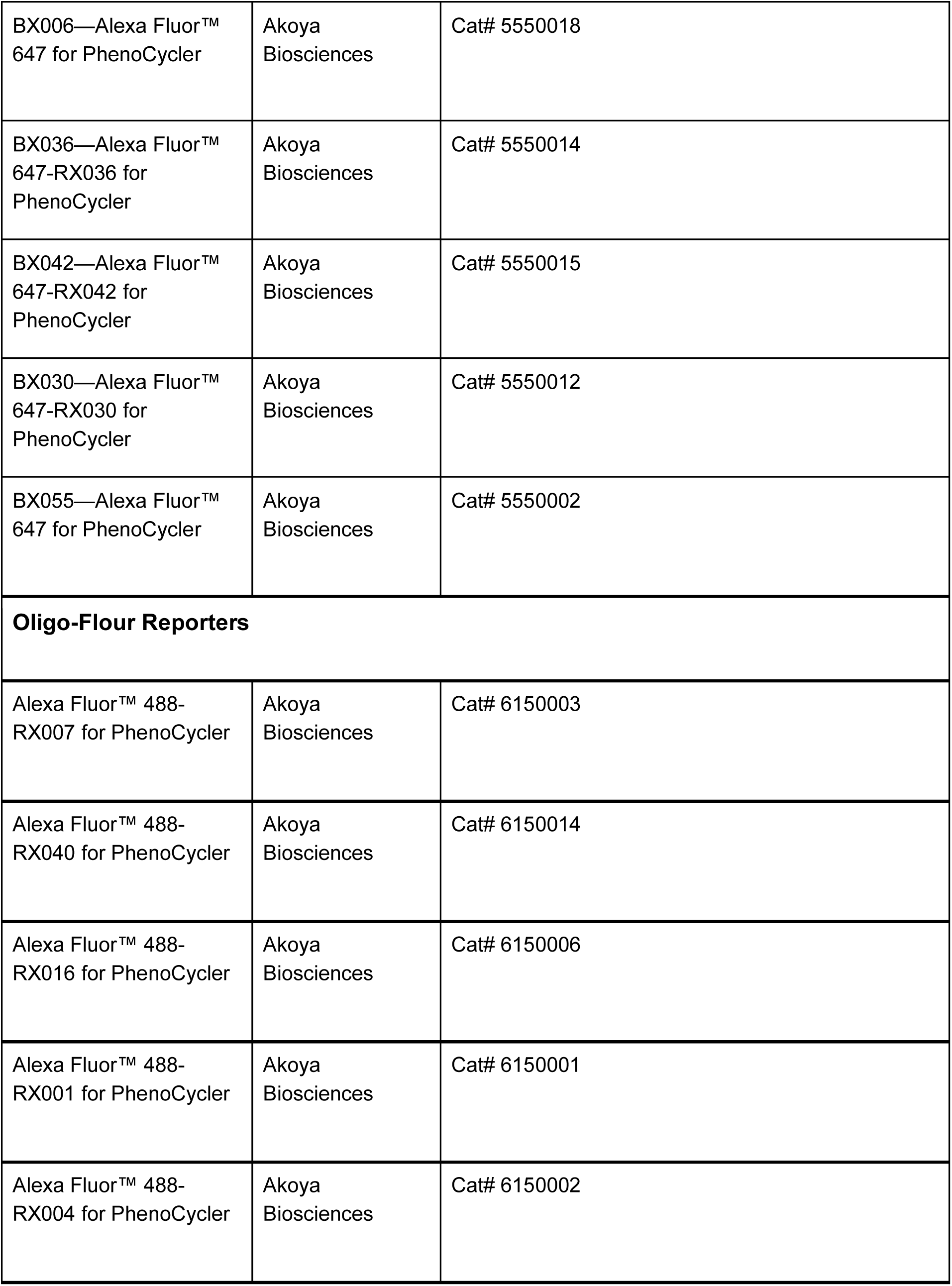
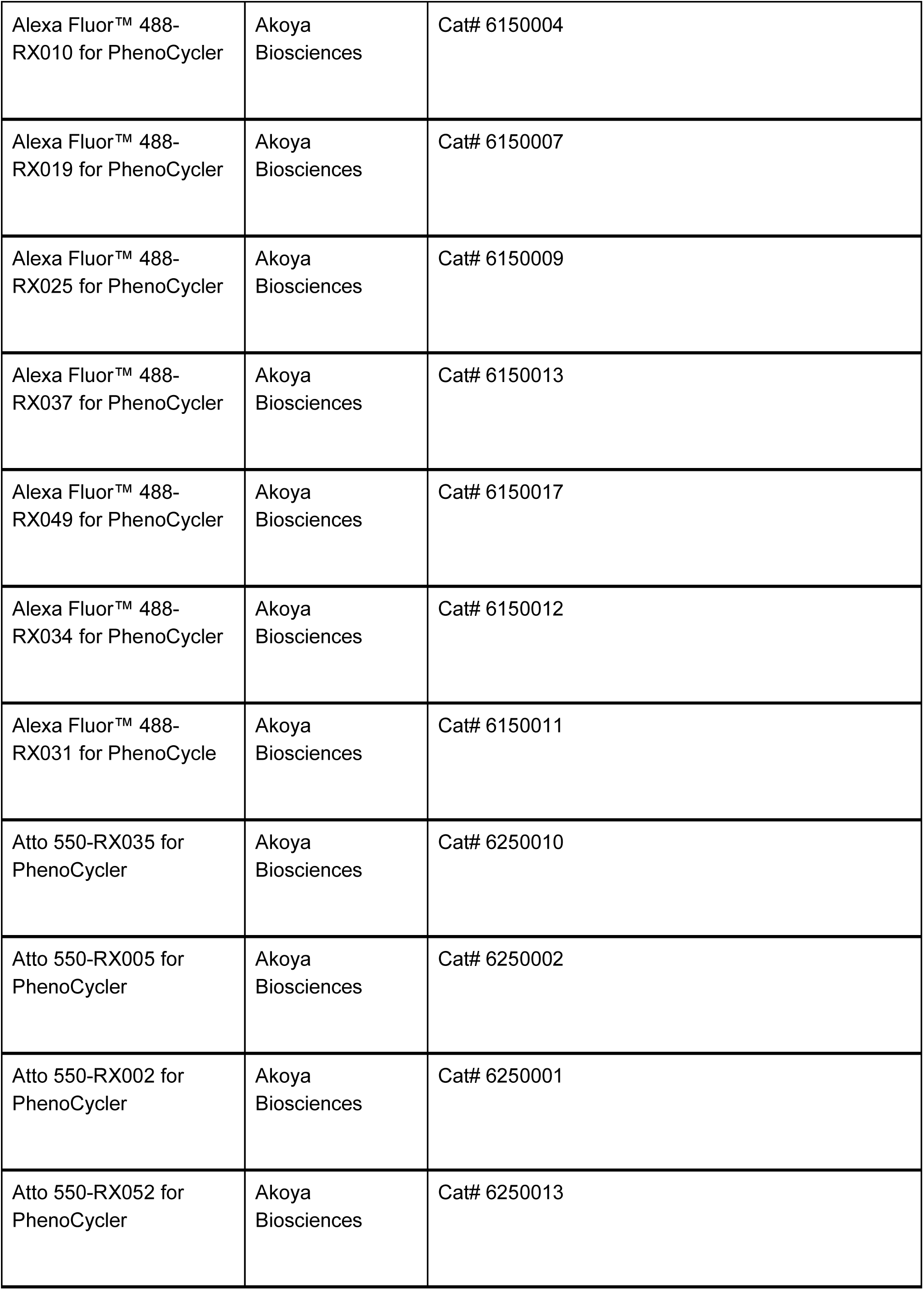
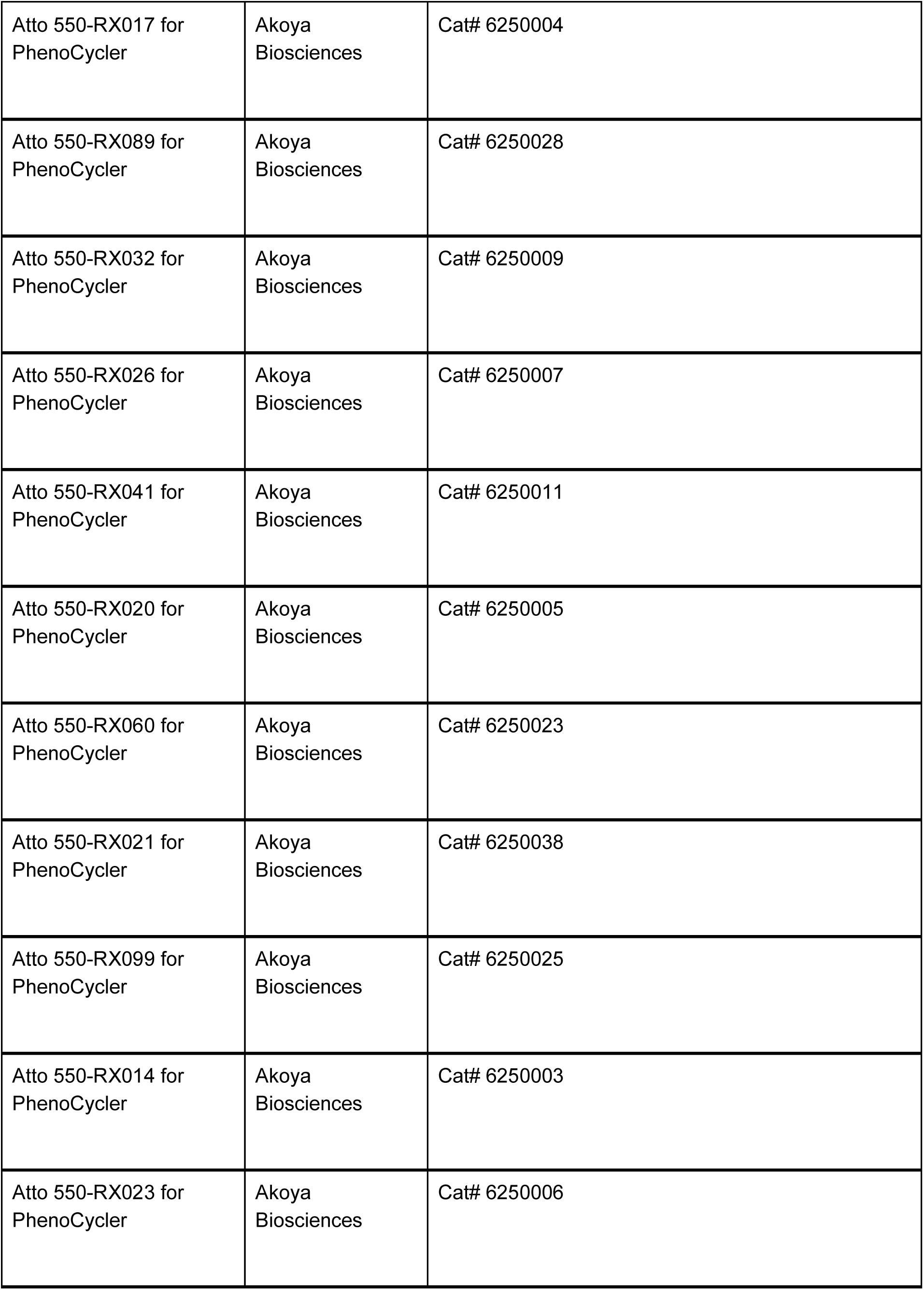
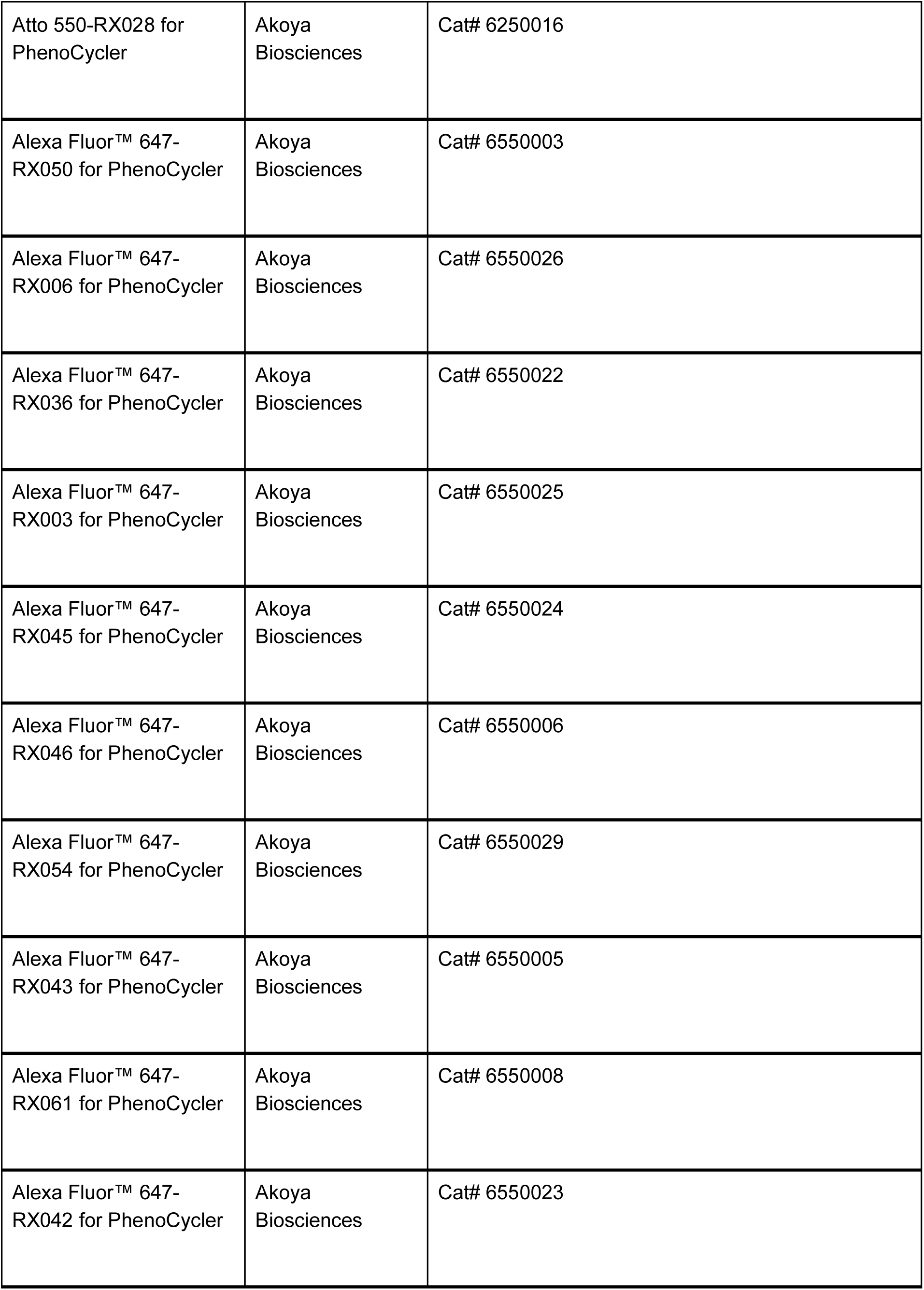
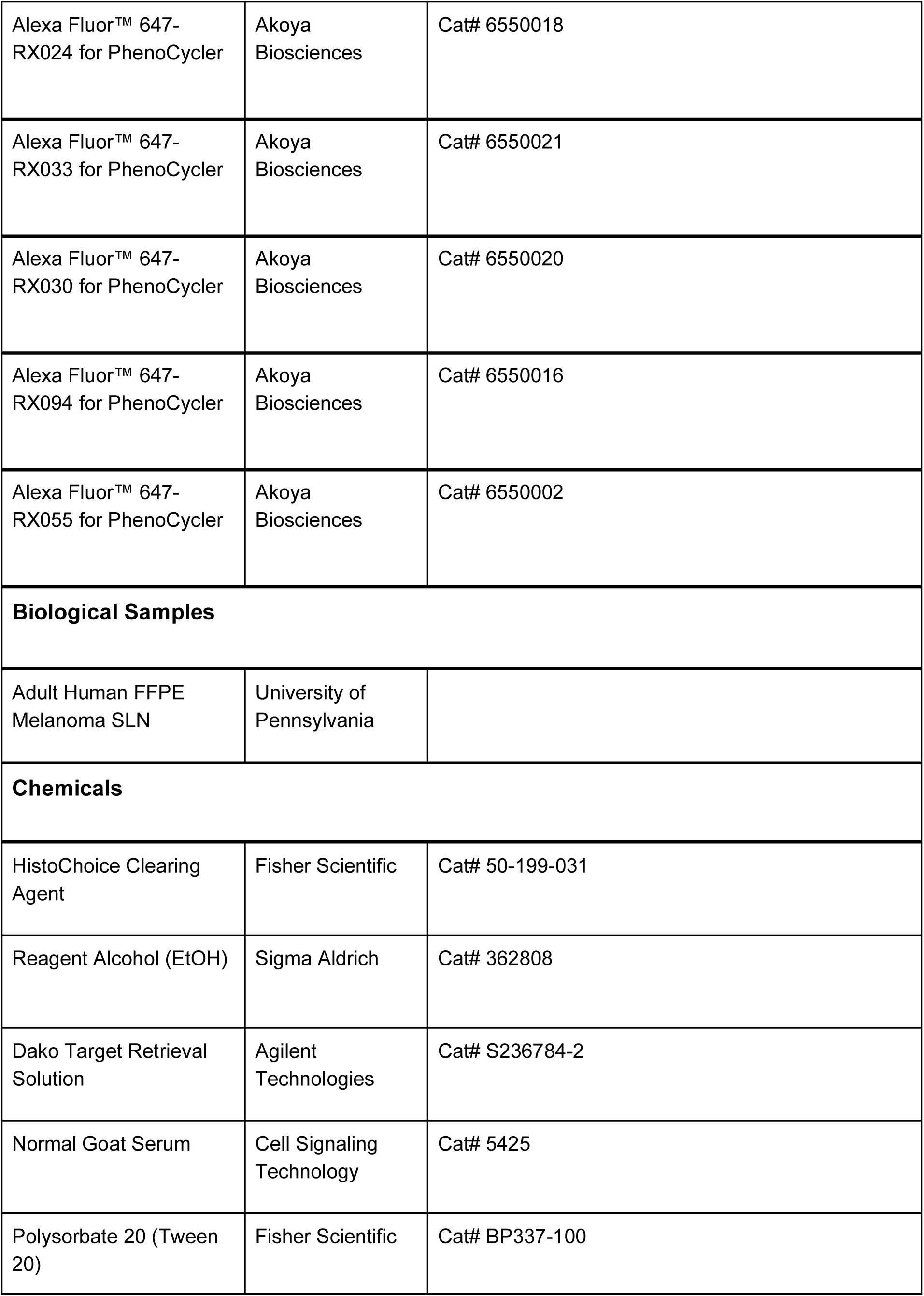
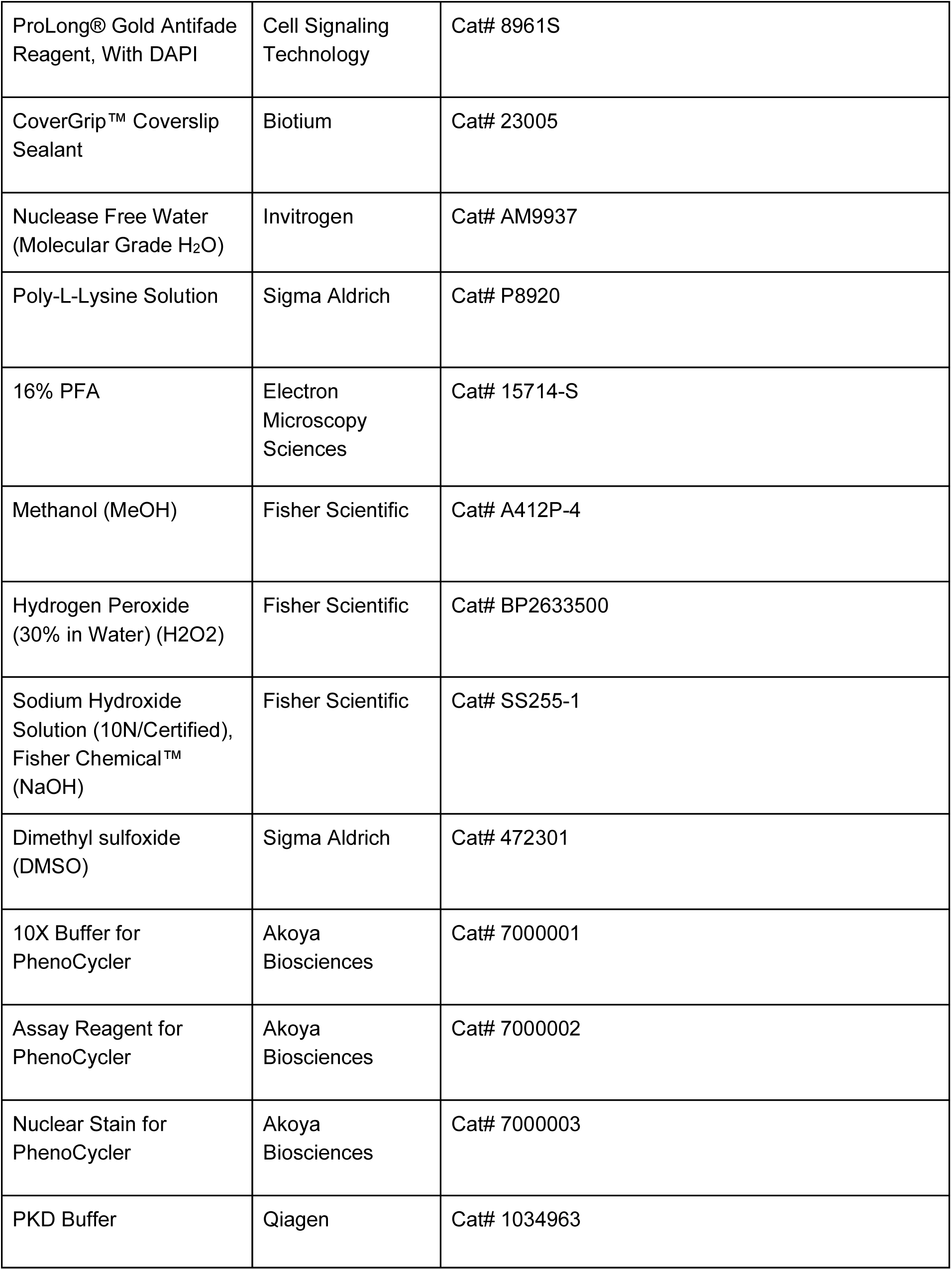
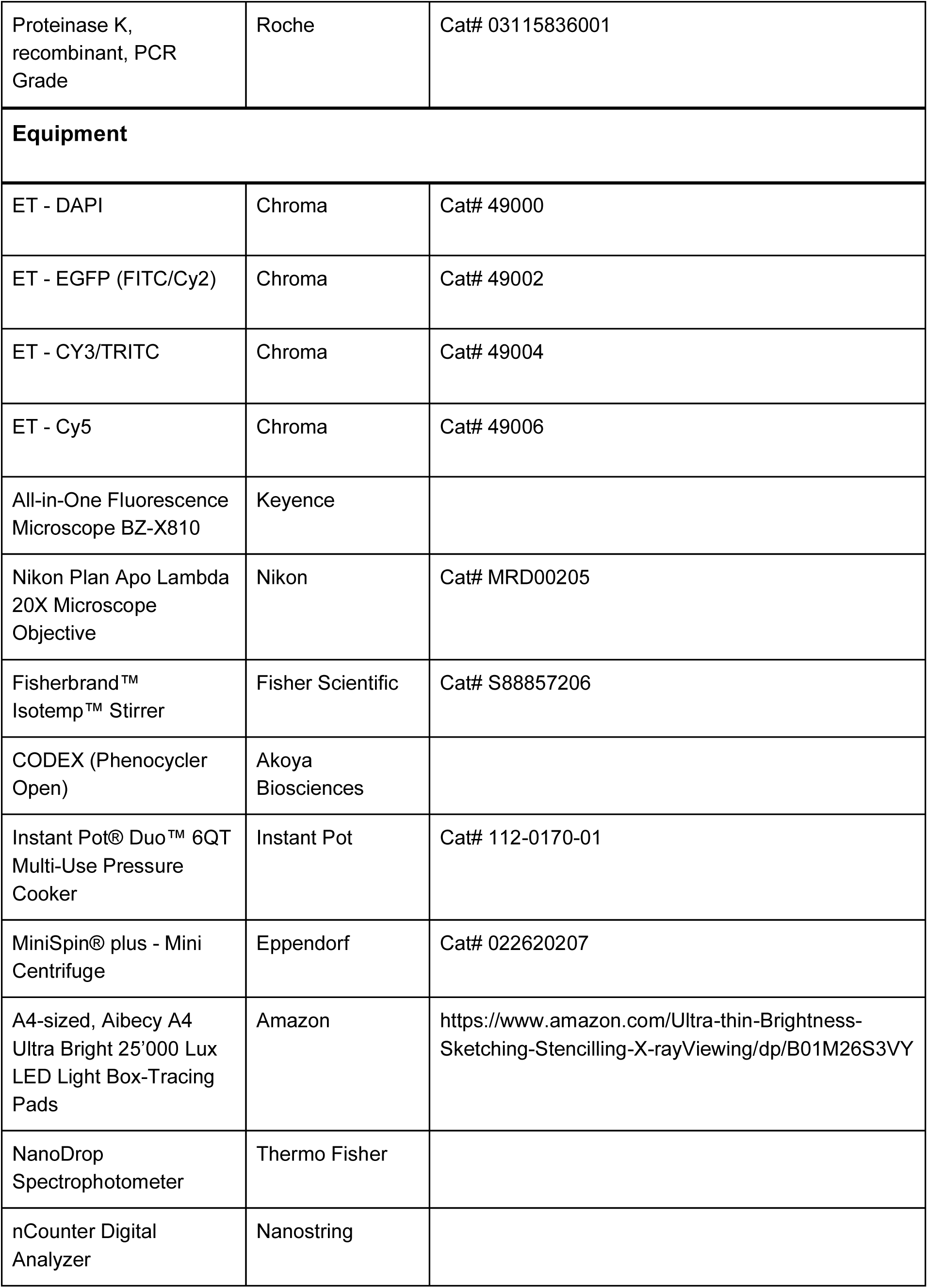
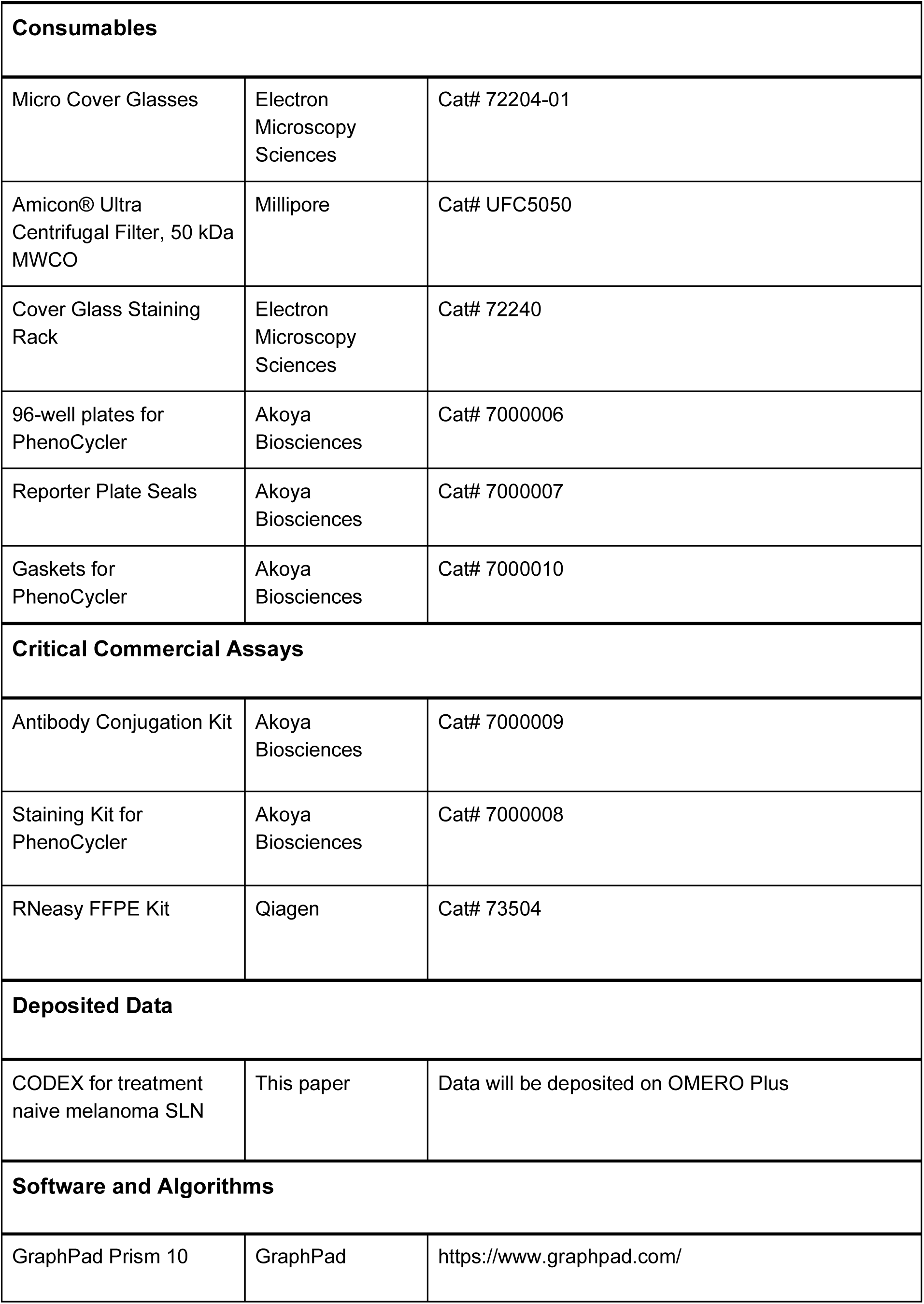
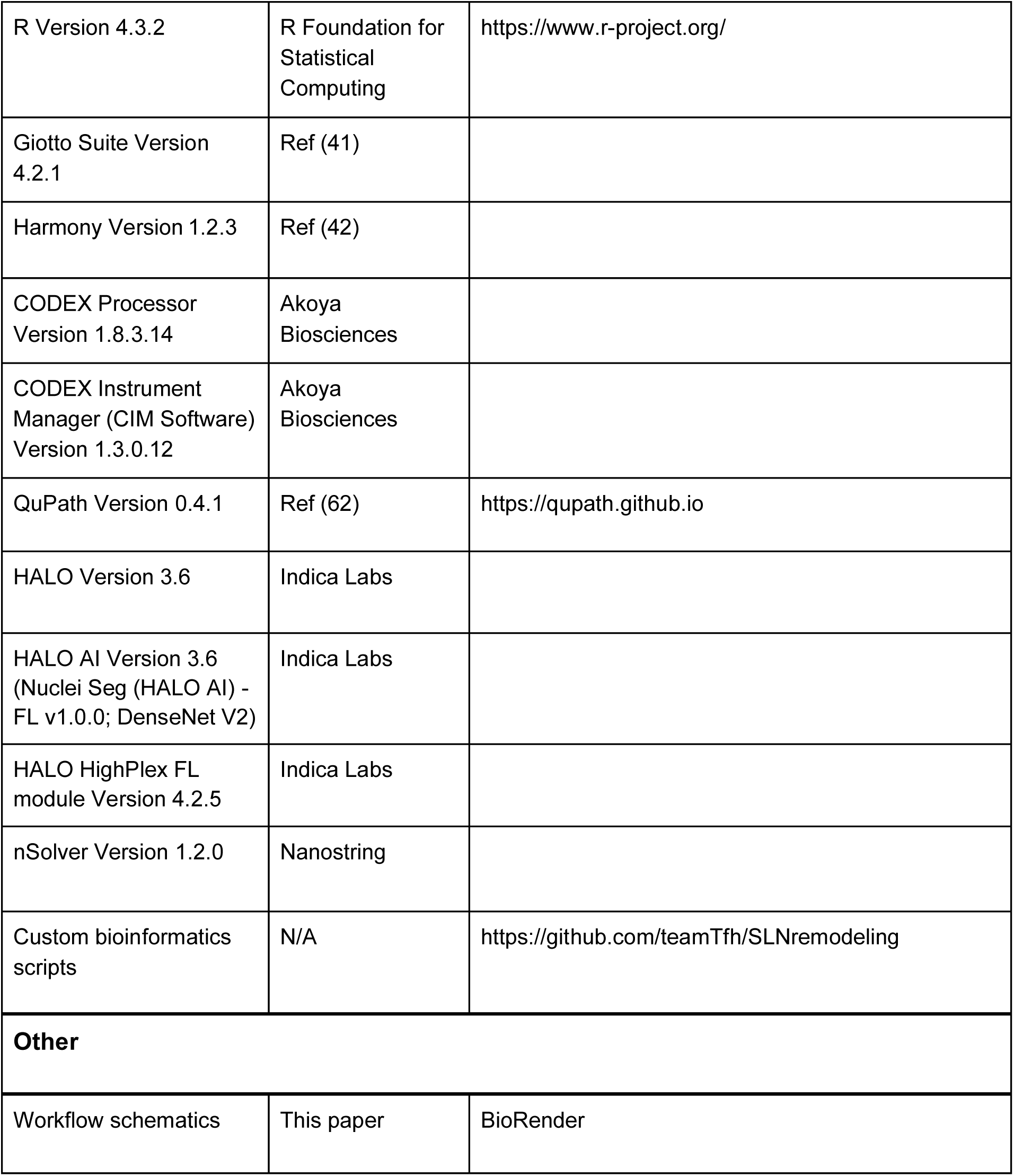

## Acknowledgements

We thank the Genomics Shared Resource at the Wistar Institute. We also thank Dr. Cindy Loomis and the staff of the Experimental Pathology Research Laboratory for their helpful guidance, as well as the Center for Biospecimen Research and Development at NYULH. We are grateful to Drs. Bertram Bengsch, Josephine Giles, Amy Baxter, Laura Vella, Erietta Stelekati, and Derek Oldridge for insightful comments and discussion. Finally, we would like to thank all the participants who have contributed to our studies.

## Author Contributions

Conceptualization: S.S., A.C.H. and R.S.H.

Data curation: S.S., A.D.

Formal analysis: S.S., Y.Y., Y.H.F., R.S.H.

Funding acquisition: R.S.H. and A.C.H.

Investigation: S.S., A.D., S.L.G-G.

Methodology: S.S., R.S.H., A.C.H.

Project administration: R.S.H.

Supervision: R.S.H.

Validation: S.S., R.S.H.

Visualization: S.S., R.S.H.

Writing – original draft: S.S., A.C.H. and R.S.H.

Writing – review & editing: S.S., Y.Y., Y.H.F., A.D., K.M., M.F., S.R., S.L.G-G., G.K., G.X., R.V., A.C.H. and R.S.H.

## FIGURE LEGENDS

**Supplemental Figure 1.**
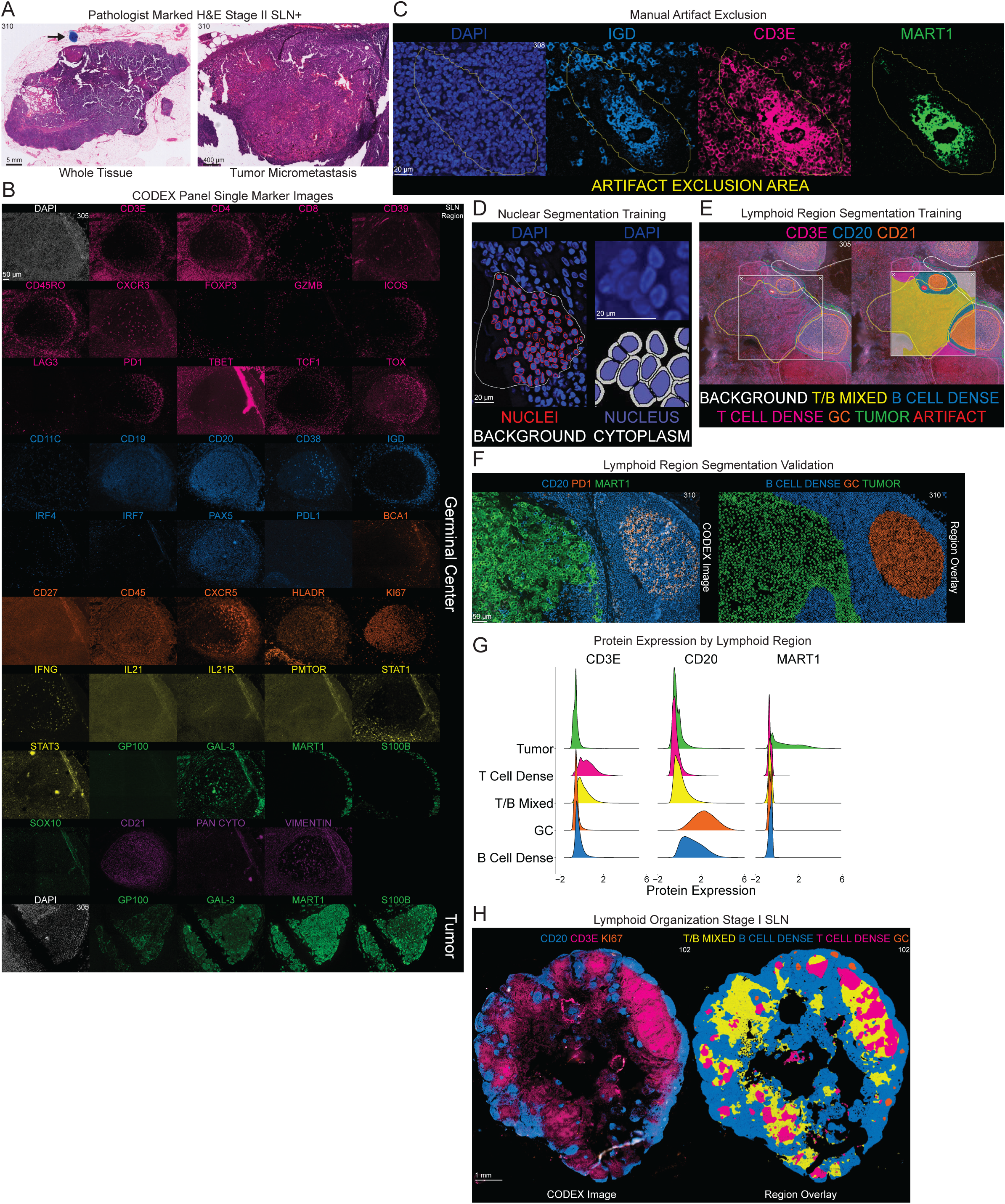
Multiplexed Immunofluorescence Imaging of the Melanoma SLN. **A.** Pathologist-marked H&E of Stage II Melanoma SLN for whole tissue (left) and tumor micrometastasis (right). Blue dot identifies tumor micrometastasis. **B.** Single marker images from Melanoma SLN tissue with either Germinal Center or Tumor field of view. **C.** Example tissue region manually marked as artifact for exclusion from downstream analysis. Image displays artifact costaining of IGD (blue), CD3E (pink), MART1 (green) and manual artifact annotation (yellow). **D.** Example tissue region manually annotated for training of Nuclear Segmentation algorithm (left) and representative image of nuclear and cytoplasmic cellular mask (right). Boundaries of nuclei in red, non-nuclei containing tissue (background) in white, and DAPI marked in royal blue (left). Nuclear mask in blue and cytoplasmic mask in white (right). **E.** Example tissue region manually annotated for training of Lymphoid Region Segmentation algorithm. Region annotated for Background (white), T/B Mixed (yellow), B Cell Dense (blue), T Cell Dense (pink), Germinal Center (orange), Tumor (green) and Artifact (red). HALO RTT window overlayed to display performance of algorithm. **F.** CODEX image of Stage II SLN (left) marking CD20 (blue), CD3E (pink), KI67 (orange), and MART1 (green) and corresponding region overlay (right) marking B Cell Dense (blue), Germinal Center (orange), and Tumor (green) regions. **G.** Protein expression of CD3E, CD20, and MART1 by lymphoid region. **H.** Stage I SLN (left) marking CD20 (blue), CD3E (pink), and KI67 (orange) and corresponding region overlay (right) marking T/B Mixed (yellow), B Cell Dense (blue), T Cell Dense (pink), and Germinal Center (orange) regions.

**Supplemental Figure 2.**
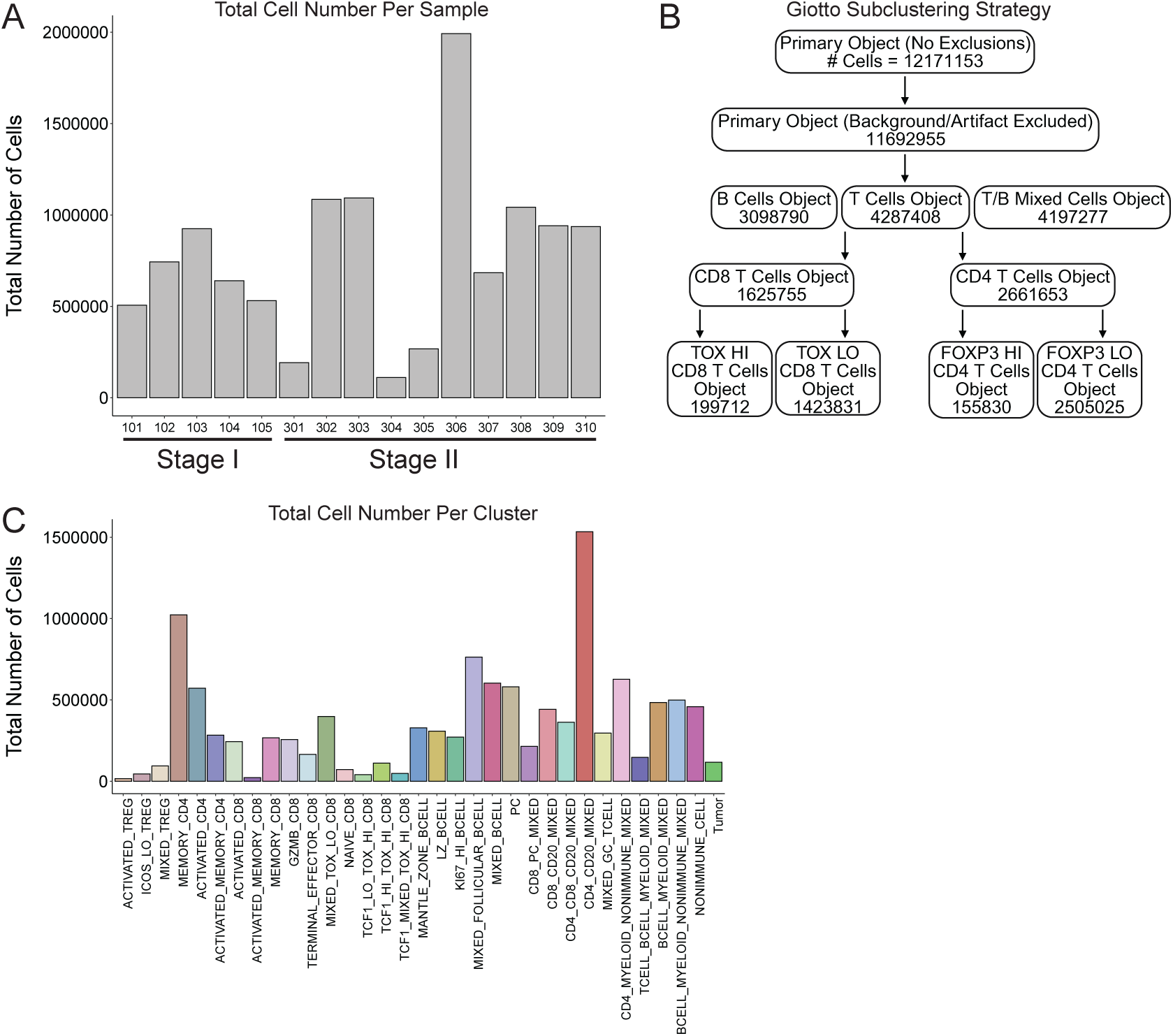
Generating a Cellular Atlas of the Melanoma SLN. **A.** Total number of cells per SLN sample. **B.** Schematic for subclustering strategy of Giotto object. Number indicates total number of cells within each subobject. **C.** Total number of cells per cluster.

**Supplemental Figure 3.**
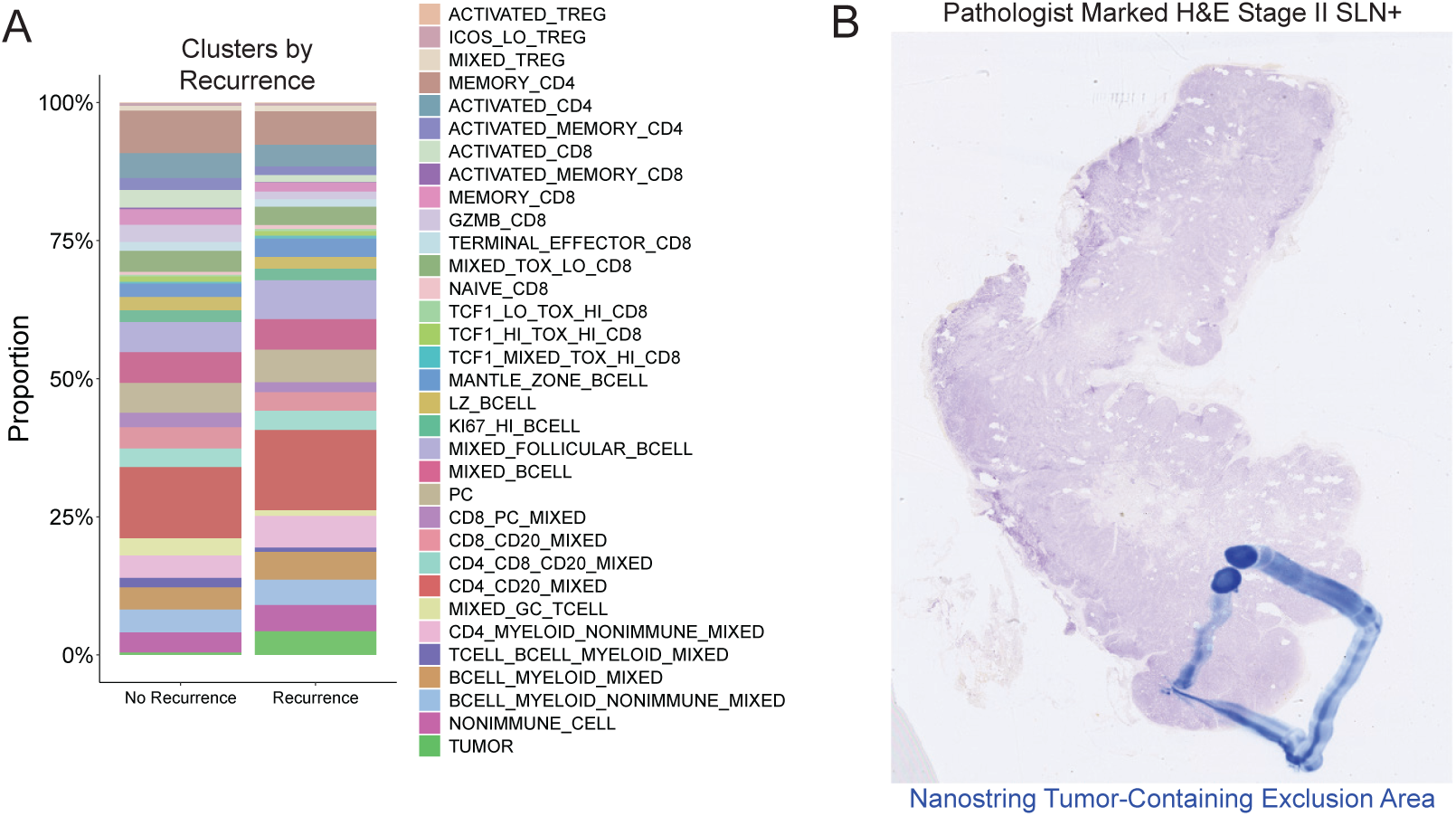
Immuno-Spatial Characterization Reveals T Cell Alterations with Stage II SLN. **A.** Cluster frequencies averaged across Stage II SLN recurrence outcome. **B.** Pathologist-marked H&E of Stage II Melanoma SLN. Blue square identifies tissue region with tumor micrometastasis excluded from Nanostring analysis.

**Supplemental Figure 4.**
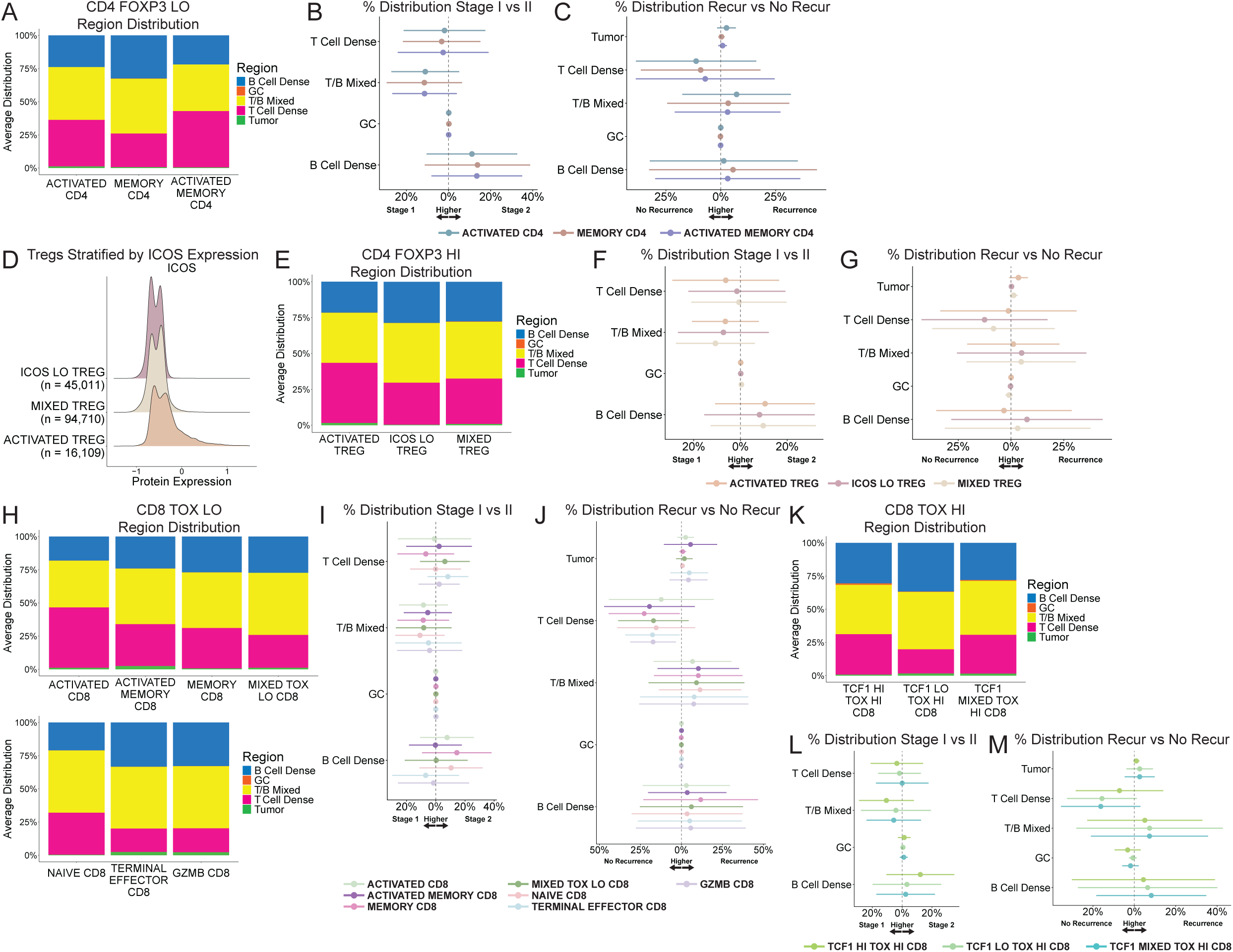
T Cells Distributed in Immune Active Regions in Melanoma SLN. **A.** ACTIVATED CD4, MEMORY CD4 and ACTIVATED MEMORY CD4 T cell distribution by region. **B-C**; **F-G**, **I-J**, **L-M.** Forest plots displaying the mean difference in proportion of cells localizing to lymphoid regions across Stage (B,F,I,L) and (C,G,J,M) Stage II Recurrence Outcomes. Circles represent the mean difference and error bars represent the 95% confidence interval. Error bars that do not intersect the vertical 0% dotted line are statistically significant. **D.** Protein expression of ICOS by ACTIVATED TREG, ICOS LO TREG and MIXED TREG populations. **E.** ACTIVATED TREG, ICOS LO TREG and MIXED TREG T cell distribution by region. **H.** ACTIVATED CD8, ACTIVATED MEMORY CD8, MEMORY CD8, MIXED TOX LO CD8, NAIVE CD8, TERMINAL EFFECTOR CD8, and GZMB CD8 T cell distribution by region. **K.** TCF1 HI TOX HI CD8, TCF1 LO TOX HI CD8, and TCF1 MIXED TOX HI CD8 T cell distribution by region.

**Supplemental Figure 5.**
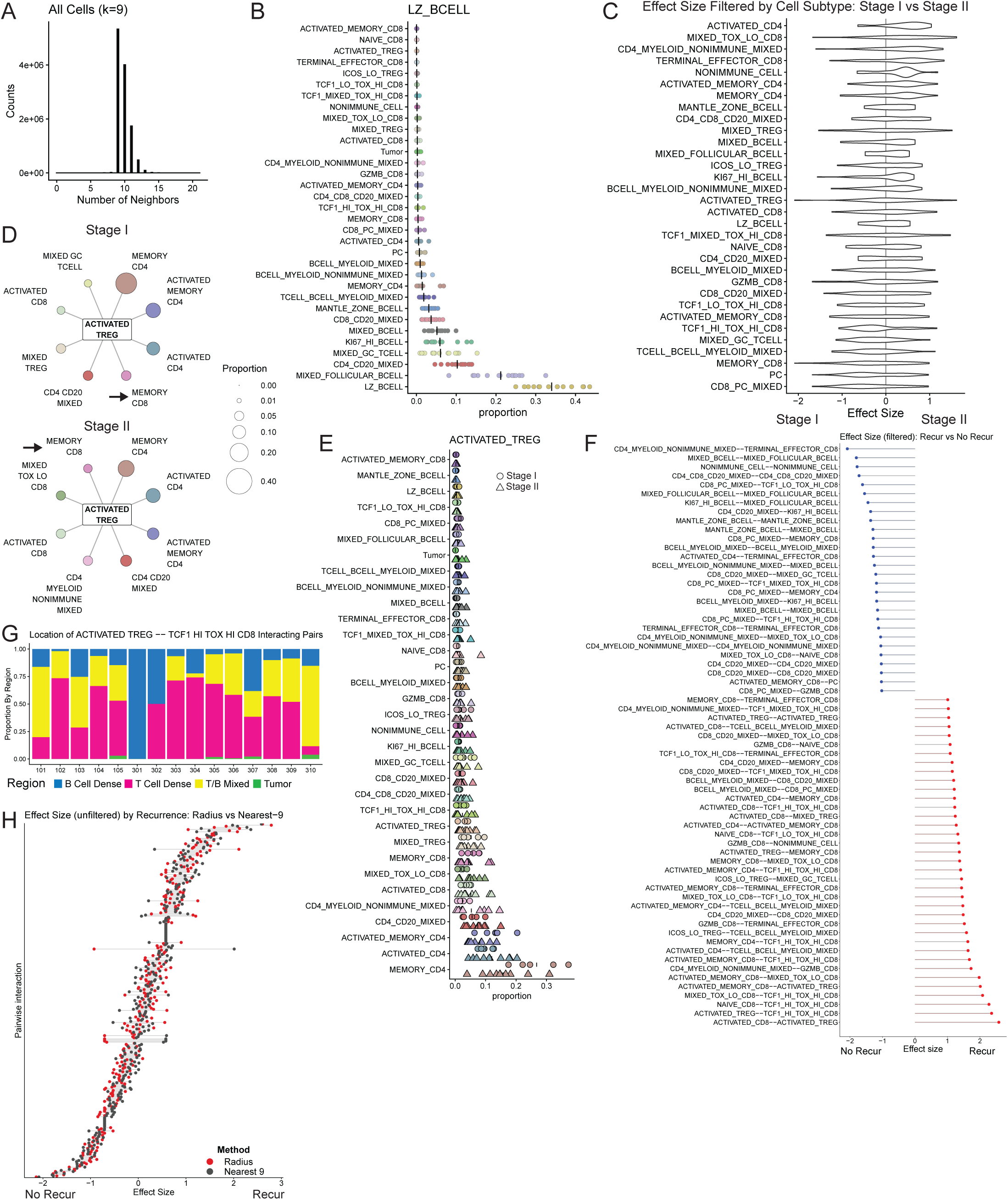
Activated Tregs Interact with Exhausted CD8 T Cells in Stage II Recurrence SLN. **A.** Histogram of number of neighbors for index cell for all cell subtypes when k=9. **B.** Dot plot displaying neighbors of LZ BCELLS by proportion of all LZ BCELL neighbors. **C.** Effect size summary plot of cell subtypes involved in interaction pairs enriched in Stage I vs Stage II SLN. **D.** Spoke plot displaying top 8 most frequent neighbors of ACTIVATED TREGS by proportion of all ACTIVATED TREG neighbors across stage. **E.** Dot plot displaying neighbors of ACTIVATED TREGS by proportion of all ACTIVATED TREG neighbors, stratified by stage (Stage 1 = circle, Stage II = triangle). **F.** Effect size of interaction pairs enriched in Stage II Recur (red) vs No Recur (blue) SLN. Effect size calculated as the difference in the mean Log2FC enrichment of interaction pairs relative to random relative to the pooled standard deviation. **G.** Stacked bar graph displaying ACTIVATED TREG -- TCF1 HI TOX HI CD8 interacting pairs distribution by region per sample. **H.** Effect size of interaction pairs enriched in Stage II Recur vs No Recur SLN, comparing the effect sizes obtained for interacting pairs identified by Nearest Neighbor (grey) or Radius (red) approach.

**Supplemental Figure 6.**
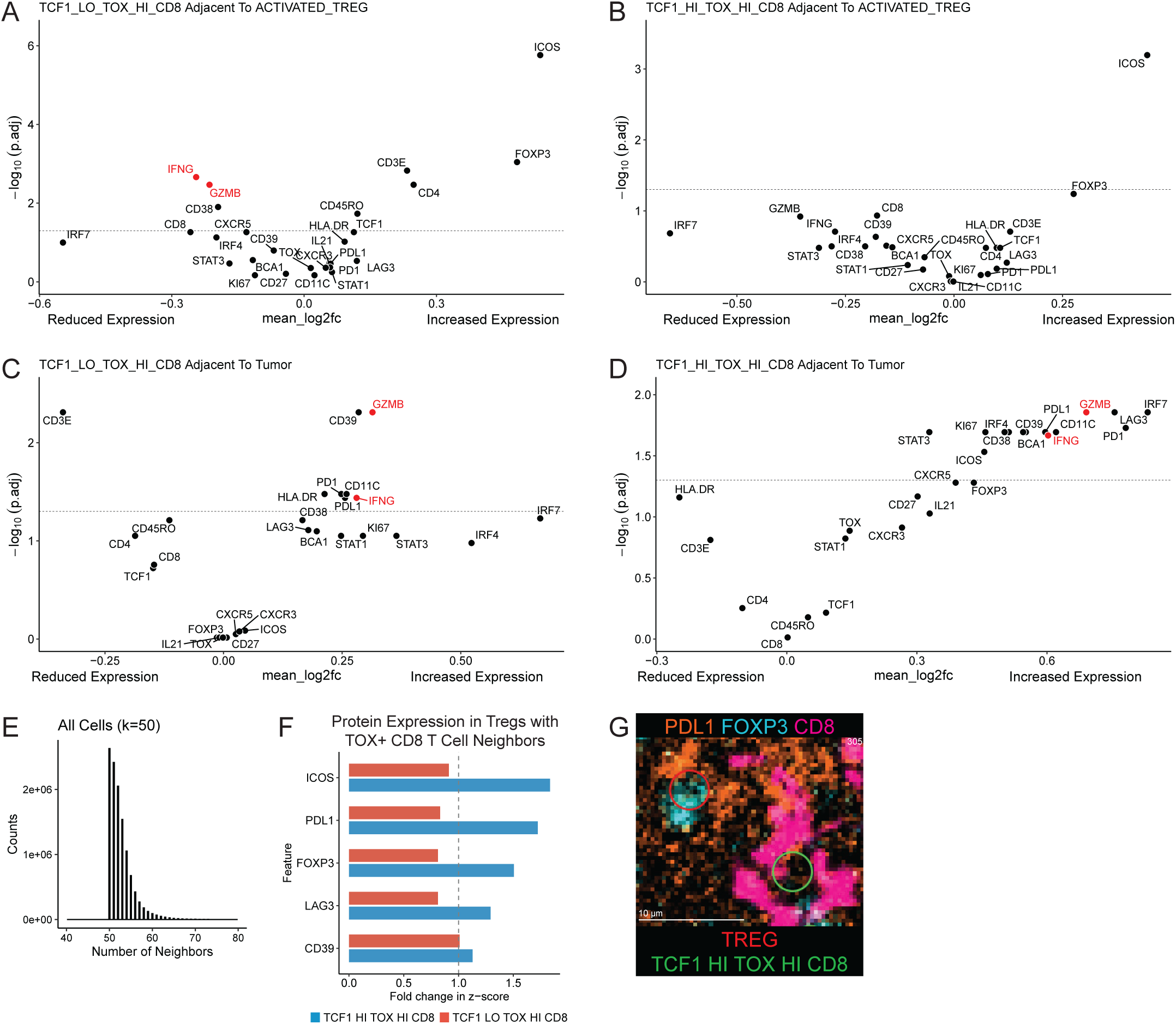
Proximal Tregs Influence Protein Expression of Exhausted CD8 T Cells in Melanoma SLN. **A-B.** Volcano plots displaying enrichment of protein expression in (A) TCF1 LO TOX HI CD8 or (B) TCF1 HI TOX HI CD8 adjacent to Activated Tregs compared to subtype interactions with all other non-Activated Treg cells. Dotted line at y=1.3 corresponding to P.adj=0.05. **C-D.** Volcano plots displaying enrichment of protein expression in (C) TCF1 LO TOX HI CD8 or (D) TCF1 HI TOX HI CD8 adjacent to Tumor cells compared to subtype interactions with all other non-Tumor cells. Dotted line at y=1.3 corresponding to P.adj=0.05. **E.** Histogram of number of neighbors across index cell - nearest neighbors relationships for all cell subtypes when k=50. **F.** Bar plot displaying fold-change in select protein expression within Tregs with TCF1 HI TOX HI CD8 T cell neighbors (blue) or TCF1 LO TOX HI CD8 T cell neighbors (red) compared to Tregs not involved in interaction. Bars to the right of the grey dotted line have a fold-change > 1.0 for proteins enriched when Tregs are involved in TOX+ CD8 T cell interactions. **G.** Example CODEX image of TREG -- TCF1 HI TOX HI CD8 interacting pairs; red circle indicates ACTIVATED TREG defined by FOXP3 (cyan), green circle indicates TCF1 HI TOX HI CD8 defined by CD8 (dark pink). Image shows PD-L1 (orange) in the Treg subset involved in TCF1+ TOX+ CD8 T cell interaction.

